# A critical role for brain nutrition in the life-history decisions of a partially migratory fish

**DOI:** 10.1101/2025.06.19.660559

**Authors:** J. Peter Koene, Arne Jacobs, Libor Závorka, Matthias Pilecky, Hannele M. Honkannen, Martin J. Kainz, Kathryn R. Elmer, Colin E. Adams

## Abstract

The role played by omega-3 long-chain polyunsaturated fatty acids (n-3 LC-PUFA) in life-history polymorphisms in partially migratory species remains poorly understood. Yet, brain development is highly dependent upon nutrition, particularly the supply of n-3 LC-PUFA, derived from diet or internally converted from their shorter-chain precursors, and the fitness of animals may be shaped by cognitive performance, including effective spatial navigation required by migration. We investigated juveniles of a wild polymorphic population of brown trout, *Salmo trutta*, with three distinct migratory ecotypes, at the point of first outward migration. Using a combination of fatty acid contents, compound-specific stable isotope analysis, and liver transcriptomics, we found that non-migrants compensated for dietary deficiency by biosynthesising n-3 LC-PUFA from precursor molecules and routing them to cell membranes to a greater extent than did migrants. These findings highlight contrasting intake and processing between migratory and non-migratory life histories of nutrients associated with brain development.

## Introduction

Migration is a common occurrence and has evolved repeatedly across the spectrum of animal phyla (*1*). This life-history strategy allows individuals to exploit the environmental heterogenies among separate habitats (*2*), yet it has costs in terms of energy expenditure and increased risk of mortality (*3*). Many species exhibit partial migration, in which migrating and non-migrating individuals occur within the same breeding population (*4*). Despite extensive research, the drivers behind the migratory decision remain uncertain (*4*). For many species, there may be strong family-level predispositions to migrate or remain resident, which are not apparent at the population level (*5*). Genetic inheritance is not decisive, however, as it has been shown that even siblings may employ different strategies (*5*). Whether to migrate may depend, in part, on physiological condition; *e.g.*, energetic state, metabolic rate, lipid storage, etc., influenced by biotic and abiotic environmental factors (*5*). Condition of potential migrants may be limited, in turn, by the quality and quantity of food resources in nursery habitats (*4*).

Animals rely on their cognitive performance for effective spatial navigation and migration, as well as foraging, finding mates, and predator avoidance (*6–9*). Brain development is highly dependent upon nutrition across many taxa, particularly the amount and type of omega-3 long-chain polyunsaturated fatty acids (n-3 LC-PUFA) integrated into neuronal membranes (*10*, *11*). These PUFA, especially docosahexaenoic acid (DHA 22:6n-3), are important for neural development and function (*12*), increasing cellular membrane fluidity and facilitating signal transfer in neural tissues due to their complex three-dimensional shapes which resist tight packing together (*10*).

Between nursery and more productive foraging habitats used at maturity, individuals may often experience environments that differ in their availability of dietary n-3 LC-PUFA. For partially migratory fluvial species, such as many salmonids and lampreys, gradients exist from n-3 LC-PUFA-poor headwaters (*13*), where juveniles undergo early development (*14*), to n-3 LC-PUFA-rich lacustrine (*15*) or even richer marine habitats (*12*, *14*). We speculate that such a gradient may provide motivation for migration. However, spending early life in an environment depauperate in n-3 LC-PUFA may itself act as a barrier to migration, since individuals may be unable to acquire sufficient dietary n-3 LC-PUFA for brain development needed for the sophisticated spatial navigation required by migration.

The capacities of consumers to convert shorter-chain n-3 PUFA to n-3 LC-PUFA differ between species and individuals (*16*). Conversion is the result of multiple synthetic pathways associated with genes coding for fatty acid desaturases (*Fads*) and elongases (*Elovl*) (*17*) (Fig. 1). It occurs primarily in liver cells (hepatocytes), from which n-3 LC-PUFA, especially DHA, may be distributed to the brain (*18*, *19*). As components of polar lipids (PL, mainly phospholipids), n-3 LC-PUFA are integral constituents of cell membranes; additionally, they are stored in neutral lipids (NL, mainly triacylglycerols) and activated as physiologically required (*10*, *20*). Thus, PUFA conversion, in combination with priority routing of DHA, may compensate for the lack of vital nutrients drawn from the diet. The two essential precursors for n-3 and n-6 LC-PUFA; i.e., α-linolenic acid (ALA; 18:3n-3) and linoleic acid (LIN; 18:2n-6), cannot be synthesised by animals and must be taken up by diet (*21, 22*). However, because precursors compete for the same enzymes to convert short-chain to long-chain PUFA, the n-6 pathway from dietary LIN to arachidonic acid (ARA;18:2n-6) should be investigated alongside the n-3 pathway from dietary ALA eventually to DHA to gain a fuller view of the biosynthesis of DHA (*23*) (Fig. 1). Studies of the evolutionary dynamics of n-3 LC-PUFA biosynthesis in natural systems have concentrated on comparing closely related pairs of species or populations of non-migratory fishes, and found that successful colonisation of novel habitats low in dietary n-3 LC-PUFA is associated with biosynthesis capacity (*24*, *25*). However, whether potential differences in n-3 LC-PUFA synthesis ability between ecotypes of partially migratory species play a role in determining life-history strategy has thus far not been examined.

**Fig. 1.**
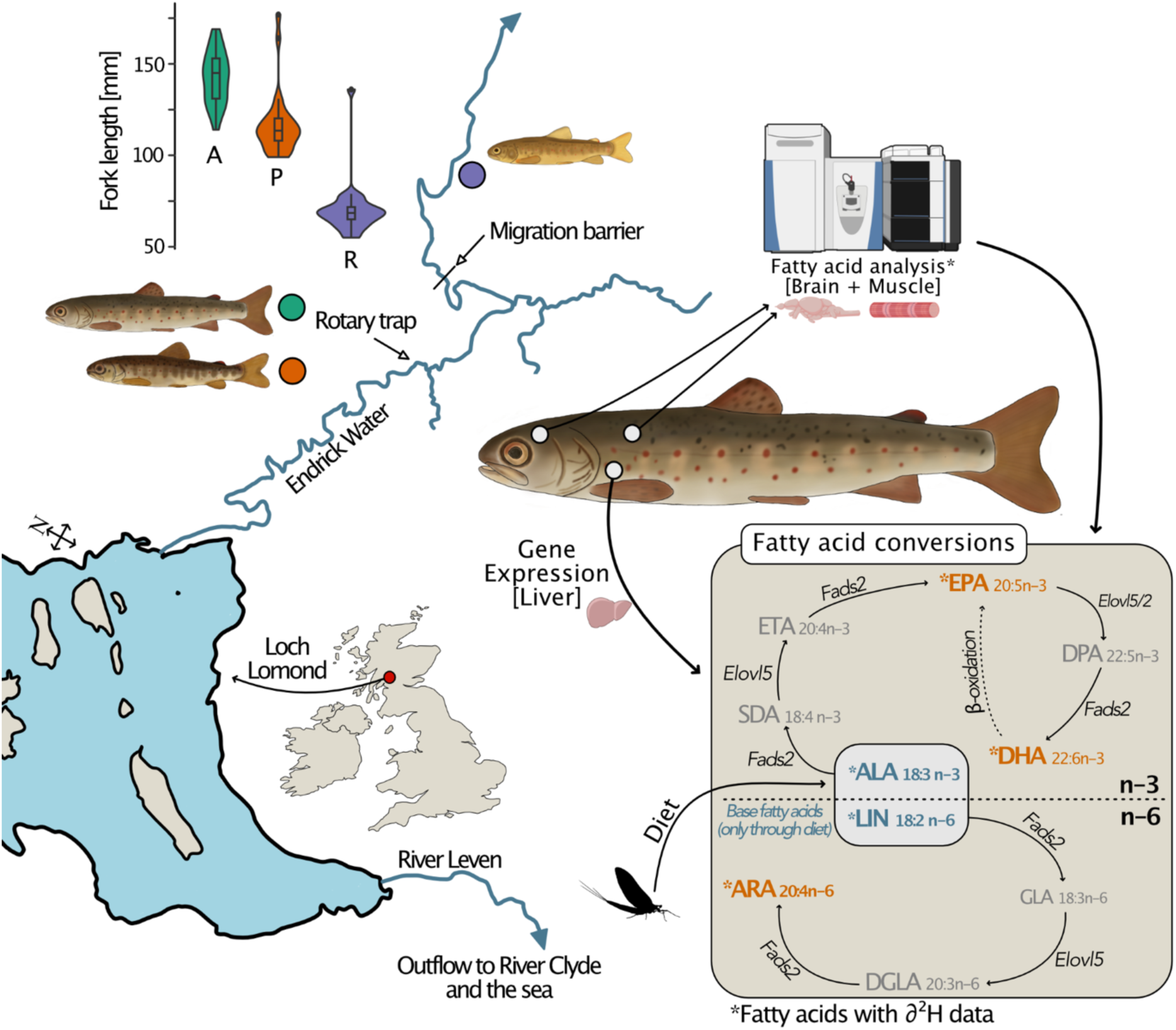
Schema-c diagram of study system. Sampling loca,ons for three migratory ecotypes of juvenile brown trout are indicated, along with comparison of ecotype sizes, n-3 and n-6 PUFA conversion pathways and associated gene expression. A = anadromous, P = potamodromous, R = riverine resident.

Brown trout (*Salmo trutta* L.) is a partially migratory, and economically and culturally important species, which is native to Eurasia and introduced world-wide. While adult and subadult forms may differ in their diets, as juveniles in their nursery streams, all trout typically have access to similar and limited food resources (*14*). Although benthic stream invertebrates on which trout feed generally lack DHA (*26*), they are typically rich in eicosapentaenoic acid (EPA; 20:5n-3), which can be subsequently converted to DHA (*27*). In contrast, terrestrial invertebrates that trout prey upon from the water’s surface tend to be richer in the short-chain n-3 PUFA, particularly in ALA (*28*), which can be converted to DHA, but at great metabolic cost (*10*, *12*). It has not yet been tested whether microhabitat foraging differences among sympatric juveniles lead to different fatty acid compositions which can be linked with different migration strategies.

We studied a wild cohort of juvenile brown trout (third spring after hatch) from a single nursery stream and composed of three migratory ecotypes on their first outward migration: anadromous (seaward migrants), potamodromous (lakeward, freshwater migrants), and riverine residents (non-migrants) (Fig. 1). We hypothesised that biosynthesis and routing of DHA are compensatory responses to dietary paucity of this nutrient, and that the degree of response associates with migratory strategy: higher capacity to biosynthesise DHA from shorter-chain precursors and subsequently route it to brain polar lipids enables a riverine resident life-history strategy, even if fulfilling energetic demands comes at the cost of somatic growth. We tested the predictions that; a) riverine residents contain lower total lipids, lower EPA and ARA contents of lipids, and different stable isotope values of ALA and LIN than migratory trout, indicating differences in diet, consistent with a greater proportion of terrestrial invertebrates; b) DHA contents are higher in brain than in muscle tissues, and higher in polar than in neutral lipids; and compound-specific stable isotopic values and gene expression would provide clear evidence of DHA biosynthesis, and; c) riverine trout biosynthesise DHA and route it to brain polar lipids to a greater extent than migratory trout, and that riverine trout show lower somatic growth than migratory trout. Finally, to test whether differential n-3 LC-PUFA biosynthesis in response to dietary paucity helps to define ecotype, we compared expression of genes directly involved in the n-3 and n-6 conversion pathways in liver tissue between the offspring of anadromous and resident brown trout, raised on the same n-3 LC-PUFA-rich diet in a common garden experiment.

## Results

### Ecotype differences in total lipids and dietary fatty acids

To determine whether ecotypes differed in their exploitation of dietary resources, we measured total lipid contents in brain and muscle tissues as well as stable hydrogen isotope (*δ*^2^H) values of the essential PUFA ALA and LIN, which trout cannot synthesise *de novo* (*21*, *22*). We further examined the percentage of lipids composed of ALA, LIN, EPA and ARA, which may indicate a terrestrial or aquatic diet source (*28*).

Ecotype had a significant effect on total lipids (Pillai = 0.879, *F*_2,16_ = 6.28, *p* < 0.001). Although an ecotype effect was not seen specifically in brain lipids, riverine trout had higher total lipid contents in dorsal muscle tissues than anadromous or potamodromous (both pairwise *post hoc* comparisons, *p* < 0.001), although there was no significant difference between anadromous and potamodromous (Fig. 2B). Compared to females, males had significantly higher total lipid contents in the brain (*F*_1,16_ = 9.38, *p* = 0.007) and muscle tissues (*F*_1,16_ = 7.85, *p* = 0.013).

**Fig. 2.**
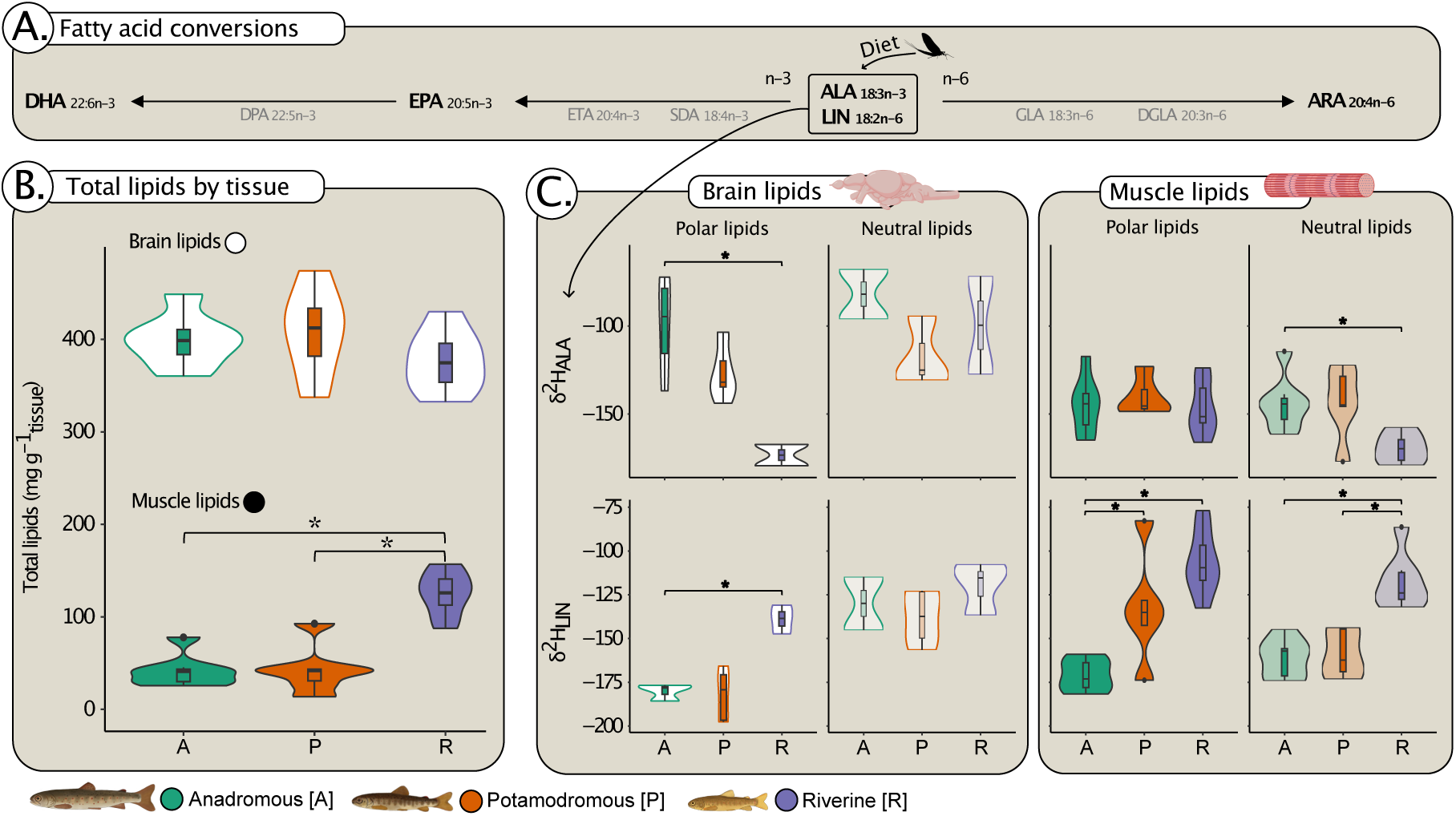
Indicators of diet differences between three migratory ecotypes of brown trout. **A.** FaFy acid conversion pathways from precursor ALA (18:3n-3) and LIN (18:2n-6); **B.** total lipids in brain and muscle, tissue; and **C.** isotopic deple,on of stable hydrogen isotopes (δ^2^H) of ALA and LIN in four lipid/,ssue types. Asterisks (*) indicate significant differences between ecotypes.

Ecotypes differed in the individual isotopic *δ*^2^H values of ALA and LIN, which are characteristic of differences in dietary intake. Values of *δ*^2^H_ALA_ differed by ecotype in brain polar lipids (PL) (*F*_2,7_ = 7.05, *p* = 0.021) and muscle neutral lipids (NL) (*F*_2,16_ = 4.7, *p* = 0.025), while *δ*^2^H_LIN_ values differed by ecotype in brain PL (*F*_2,7_ = 14.2, *p* = 0.003), muscle PL (*F*_2,15_ = 16.2, *p* < 0.001), and muscle NL (*F*_2,15_ = 21.1, *p* < 0.001) (Fig. 2C; Supplementary material: Tables S1–S4). Sex played no significant role in *δ*^2^H_ALA_ or *δ*^2^H_LIN_ values in any of the lipid/tissue types.

Riverine trout showed greater ALA contents in muscle NL than did migrants (*F*_2,16_ = 4.7, *p* = 0.025; *post-hoc*: between riverine and anadromous, and riverine and potamodromous, *p* < 0.001) (Supplementary material: Tables S5, S6). There was no difference between ecotypes in LIN contents of any lipid/tissue type (Supplementary material: Table S7). Migratory ecotypes showed higher EPA contents in brain PL than riverine residents (*F*_2,17_ = 25.66, *p* < 0.001; *post-hoc*: between riverine and anadromous, and riverine and potamodromous, *p* < 0.001) and brain NL (*F*_2,17_ = 22.64, *p* < 0.001; *post-hoc*: all pairwise comparisons, *p* < 0.001) (Supplementary material: Tables S8, S9). Migratory ecotypes also showed greater ARA contents than riverine trout in brain NL (*F*_2,17_ = 19.01, *p* < 0.001; *post-hoc*: between riverine and anadromous, *p* < 0.001, and riverine and potamodromous, *p* = 0.001) and in muscle NL (*F*_2,17_ = 4.58, *p* = 0.026; *post-hoc*: between riverine and anadromous, *p* = 0.022) (Supplementary material: Tables S10, S11).

Overall, our results show that riverine residents had higher total lipids in muscle tissues, lower EPA and higher ALA contents, and different *δ*^2^H values in essential PUFA (lower *δ*^2^H_ALA_, but higher *δ*^2^H_LIN_) than migratory trout, and are broadly consistent with our expectation of dietary reliance on terrestrial invertebrates.

### Compensatory DHA routing and biosynthesis

To discern whether different migratory ecotypes compensate for dietary lack of DHA by biosynthesis and routing of DHA to brain polar lipids, we measured DHA contents and *δ*^2^H values of DHA across polar and neutral lipids in brain and muscle tissue.

Significant differences in DHA of polar lipids in brain and muscle tissues was revealed. The DHA content of polar (membrane) and neutral (storage) lipids differed significantly in brains and muscle tissues of three brown trout ecotypes (*F*_3,60_ = 247.7, *p* < 0.001) with significantly higher DHA contents in polar lipids than in neutral lipids for brain (*post-hoc*: *p* < 0.001) and muscle tissues (*post-hoc*: *p* < 0.001). However, there was no significant difference in DHA content of brain and muscle tissues (Fig. 3; Supplementary Material: Table S12).

**Fig. 3.**
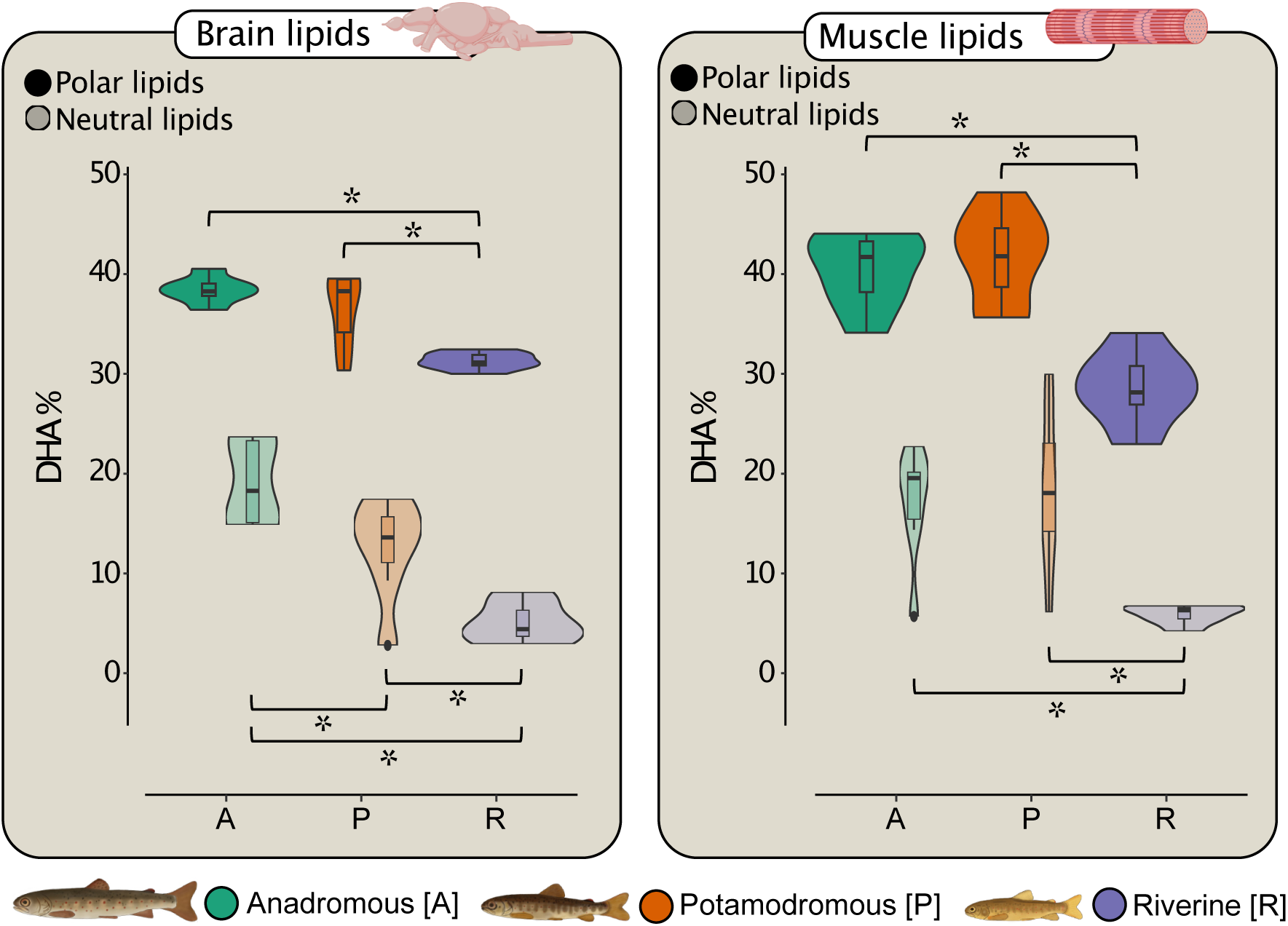
DHA contents (%) of polar (membrane) and neutral (storage) lipids in brains and muscle-ssues of three brown trout ecotypes. Asterisks (*) indicate significant differences between ecotypes.

Biosynthesis of DHA was clearly stimulated in the trout’s n-3 LC-PUFA-deprived nursery environment. Compound-specific stable isotope analysis revealed isotopic depletion of Δ*δ*^2^H_DHA_ (*i.e.* lighter hydrogen isotopes in DHA after correction for differences in individual diets) in each of the lipid classes and tissue types (mean Δ*δ*^2^H_DHA_ all < −5.7) (Fig. 4B).

**Fig. 4.**
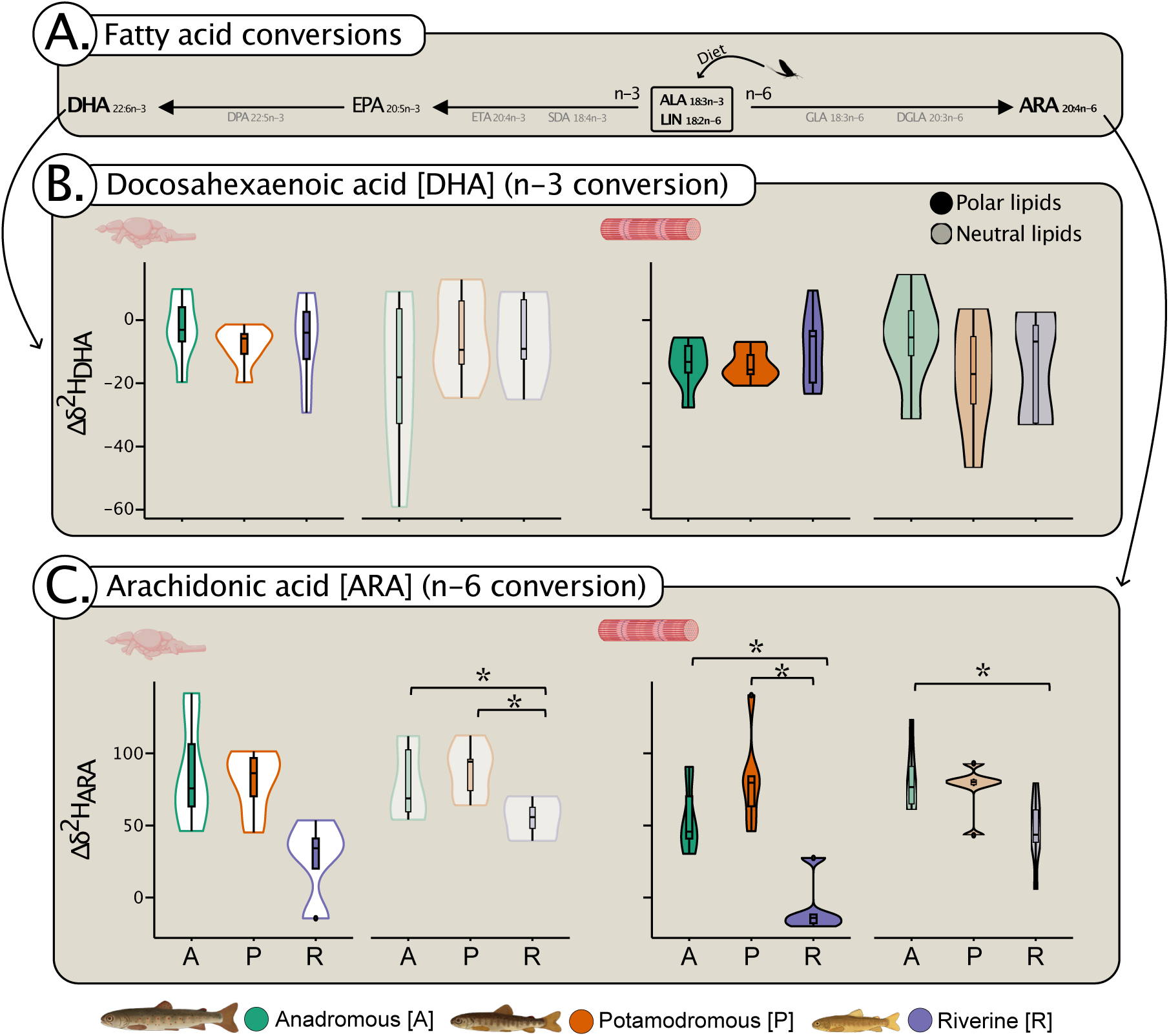
Stable hydrogen isotope values corrected for differences in diet between three brown trout ecotypes (Δ*δ*^2^H). **A.** faFy acid conversion pathways to: **B.** DHA (22:6n-3), and C. ARA (20:4n-6), across polar and neutral lipids in brain and muscle,ssue. Asterisks (*) indicate significant differences between ecotypes.

Overall, our findings show that, in a DHA-impoverished nursery environment, all ecotypes compensated for dietary DHA deficit with biosynthesis and routing of DHA to polar lipids.

### Ecotype differences in LC-PUFA contents and synthesis

Potential differences between ecotypes in DHA biosynthesis and routing were quantified to determine whether riverine trout biosynthesised DHA and routed it to brain polar lipids to a greater extent than migratory types. Depletion of hydrogen stable isotopes in the n-6 PUFA, ARA, was also measured, as n-6 LC-PUFA biosynthesis is often a corollary of n-3 LC-PUFA biosynthesis (*23*).

There were significant ecotype differences in the DHA content of lipid classes and tissue types (Table 1). Riverine trout had less DHA in every lipid class and tissue type than both migratory ecotypes. Differences between migrants showed only in brain PL, where anadromous trout had significantly greater DHA contents than potamodromous (Table 2; Fig. 3). As an indicator of biosynthesis, however, there was no significant difference between ecotypes in changes of Δ*δ*^2^H_DHA_ (Fig. 4B; Supplementary material: Table S13).

**Table 1.** ANOVA results for effects of ecotype and sex on DHA contents (%). Two lipid classes and two,ssue types (brain and muscle) were tested. PL = polar lipids, NL = neutral lipids.

| Lipid type | Variable | df | Sum Sq. | Mean Sq. | F | Pr(>F) |
| --- | --- | --- | --- | --- | --- | --- |
| Brain PL | Ecotype | 2 | 190.2 | 95.1 | 17.230 | < 0.001 |
|  | Sex | 1 | 2.0 | 2.0 | 0.370 | 0.551 |
|  | Residuals | 17 | 93.8 | 5.5 |  |  |
| Brain NL | Ecotype | 2 | 689.2 | 344.6 | 20.392 | < 0.001 |
|  | Sex | 1 | 0.1 | 0.1 | 0.008 | 0.930 |
|  | Residuals | 17 | 287.3 | 16.9 |  |  |
| Muscle PL | Ecotype | 2 | 725.6 | 362.8 | 21.355 | < 0.001 |
|  | Sex | 1 | 2.7 | 2.7 | 0.162 | 0.693 |
|  | Residuals | 17 | 288.8 | 17.0 |  |  |
| Muscle NL | Ecotype | 2 | 655.9 | 328.0 | 9.410 | 0.002 |
|  | Sex | 1 | 0.1 | 0.1 | 0.004 | 0.923 |
|  | Residuals | 17 | 592.5 | 34.9 |  |  |

**Table 2.** Pairwise comparisons from Tukey’s HSD *post-hoc* tests of the effect of ecotype on DHA contents (%). Two lipid classes and two,ssue types (brain and muscle) were tested. PL = polar lipids, NL = neutral lipids, Ana = anadromous, Pot = potamodromous, Riv = riverine resident.

| Lipid type | Comparison | Difference | Lower | Upper | $p_{adj}$ |
| --- | --- | --- | --- | --- | --- |
| Brain PL | Pot : Ana | -1.931 | -5.153 | 1.290 | 0.299 |
|  | Riv : Ana | -7.126 | -10.348 | -3.905 | < 0.001 |
|  | Riv : Pot | -5.195 | -8.416 | -1.973 | 0.002 |
| Brain NL | Pot : Ana | -6.609 | -12.246 | -0.972 | 0.021 |
|  | Riv : Ana | -14.025 | -19.662 | -8.388 | < 0.001 |
|  | Riv : Pot | -7.416 | -13.053 | -1.779 | 0.010 |
| Muscle PL | Pot : Ana | 1.341 | -4.311 | 6.993 | 0.817 |
|  | Riv : Ana | -11.745 | -17.397 | -6.093 | < 0.001 |
|  | Riv : Pot | -13.086 | -18.738 | -7.434 | < 0.001 |
| Muscle NL | Pot : Ana | 1.368 | -6.727 | 9.463 | 0.902 |
|  | Riv : Ana | -11.112 | -19.207 | -3.017 | 0.007 |
|  | Riv : Pot | -12.480 | -20.575 | -4.385 | 0.003 |

In contrast, the long-chain n-6 PUFA, ARA, differed between ecotypes in its Δ*δ*_2_H values in all lipid/tissue types, except brain NL (Fig. 4C; Supplementary material: Table S14). In each case, riverine trout showed lower values than either migratory ecotype (*post-hoc*: in brain PL compared to anadromous, *p* = 0.02; to potamodromous, *p* = 0.033; in muscle PL compared to anadromous, *p* = 0.004; to potamodromous, *p* < 0.001; and in muscle NL compared to anadromous, *p* = 0.022; but to potamodromous, *p* = 0.071), including depletion in muscle PL (Supplementary material: Table S15). However, anadromous and potamodromous did not differ from each other.

Multinomial logistic regressions tested whether each ecotype had a distinct and characteristic signature of LC-PUFA contents and isotopic depletion across lipid classes and tissue types. Combinations of DHA and EPA contents, Δ*δ*^2^H_DHA_ and Δ*δ*^2^H_EPA_, and Δ*δ*^2^H_DHA_ and Δ*δ*^2^H_ARA_ across the lipid classes and tissue types gave accurate predictions of ecotype for every specimen with probabilities of > 99.9 %. The model combining DHA and ARA contents accurately predicted the ecotype of all specimens with probabilities of > 95 %, with one exception (> 92.3 % prob.) (Supplementary material: Tables S16–S19).

In summary, riverine trout had lower DHA contents than migratory types and lower Δ*δ*^2^H_ARA_ values, indicating biosynthesis of the n-6 PUFA, ARA, whose pathway competes for the same enzymes as the n-3 pathway that leads to EPA (*23*).

### Effects on somatic growth and morphology

Potential body size differences between wild trout ecotypes were used to infer the cost to somatic growth of DHA biosynthesis, as ecotype proved to be highly colinear with the expression of *Fads2* and *Elovl5* genes known to be involved in LC-PUFA synthesis (*17*). All juveniles captured (*n* = 85) were the same age class, yet differed in size between ecotypes. Ecotype, but not sex, exerted a strong effect on fork length (*F*_2,79_ = 175.0, *p* < 0.001), with anadromous trout (142 mm ± 14 s.d.) being significantly larger than potamodromous (117 mm ± 17 s.d.), which, in turn, were significantly larger than riverine (70 mm ± 15 s.d.) (*post-hoc*: all pairwise comparisons *p* < 0.001) (Fig. 1).

### Candidate gene expression underlying fatty acid metabolism

Potential up-regulation in riverine trout of *Fads* and *Elovl* candidate genes was quantified using mRNA sequencing of the liver transcriptome. Significant divergence in gene expression across the transcriptome, generally, was found between all ecotype pairs, especially between riverine and each of the migratory types (Fig. 5A; Supplementary material: Fig. S1, Table S20). Of two differentially expressed *Fads2* loci on chromosome 4, one was up-regulated in riverine compared to both anadromous (log2 fold change, LFC = 1.257, *p* < 0.001) and potamodromous (LFC = 0.733, *p* = 0.014), while anadromous was down-regulated compared to potamodromous (LFC = −0.523, *p* = 0.079). Expression at another *Fads2* locus showed up-regulation in riverine only compared to anadromous (LFC = 0.638, p = 0.027). *Elovl2* expression was also not differentially expressed by various ecotypes, but *Elovl5* was. Of two differentially expressed *Elovl5* loci, the one on chromosome 10 was up-regulated in riverine compared to both anadromous (LFC = 1.387, *p* < 0.001) and potamodromous (LFC = 0.809, *p* = 0.008), but down-regulated in anadromous compared to potamodromous (LFC = −0.578, *p* = 0.059). Elovl5 on chromosome 18 differed significantly only between riverine and the two migratory types (riverine-anadromous: LFC = 2.753, *p* < 0.001; riverine-potamodromous: LFC = 2.827, *p* < 0.001) (Fig. 5B; Supplementary material: Table S20).

**Fig. 5.**
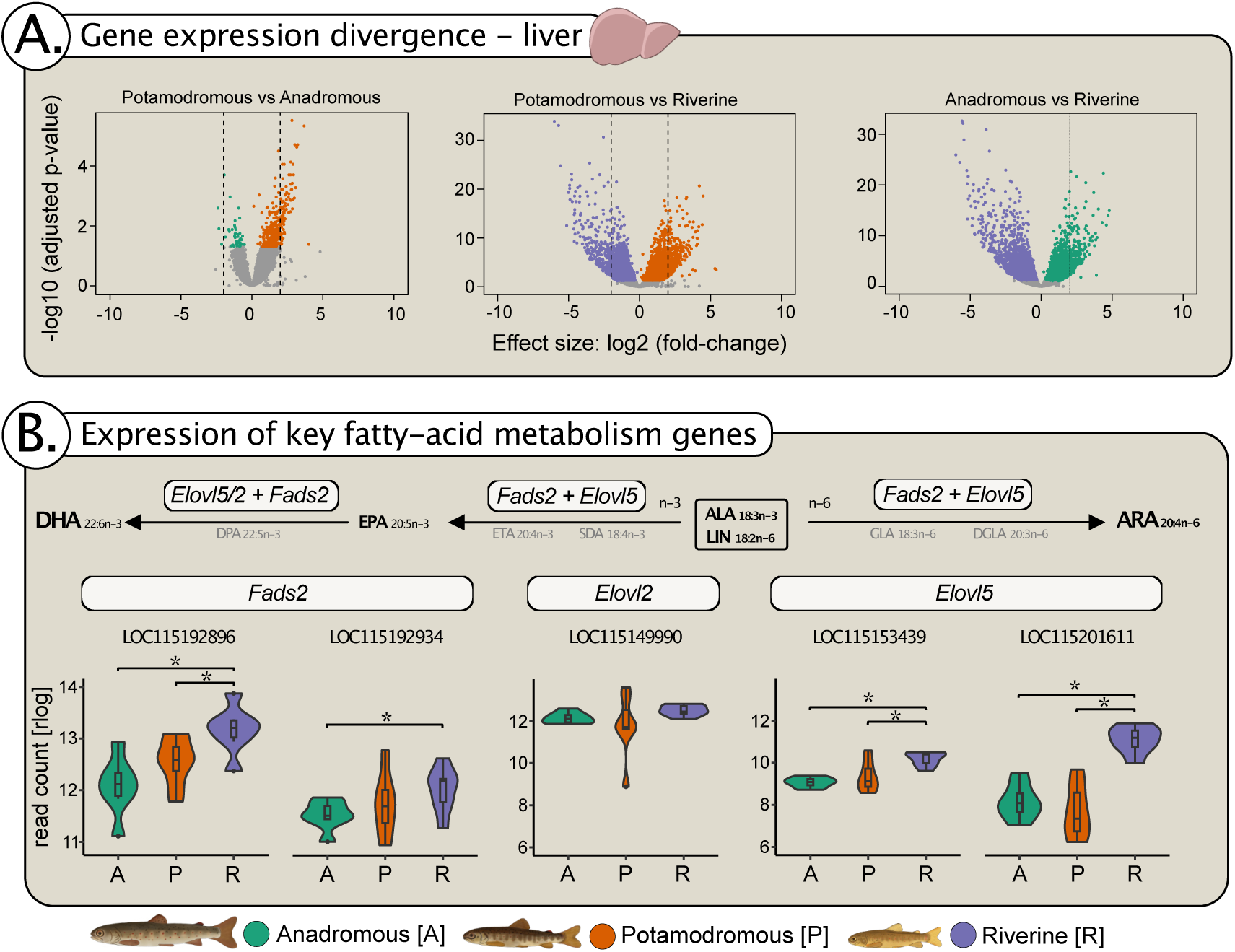
Gene expression in liver -ssue of three brown trout ecotypes. A. divergence in expression across 24,993 genes between migratory ecotypes, shown as log of adjusted pvalues on effect sizes between pairwise ecotype comparisons; B. rlog read counts from mRNA sequences of candidate genes involved in LC-PUFA biosynthesis (*Fads2* at two loci, *Elovl2*, and *Elovl5* at two loci) in three brown trout ecotypes. Asterisks (*) indicate significant differences between ecotypes.

### Co-expression and gene ontology

A weighted co-expression network analysis was conducted *a posteriori* to determine the association of key gene co-expression modules with ecotype and LC-PUFA bioconversion. This analysis revealed 28 modules of gene co-expression. Nineteen modules were significantly correlated with negative Δ*δ*_2_H values, indicating isotopic depletion, in LC-PUFA across the two lipid classes and two fish tissue types; 11 of these modules were also associated with ecotype (Fig. 6A). The arbitrarily named ‘black’ module, with 3688 genes, showed higher cumulative expression (module eigengene values) in riverine trout than either of the migratory ecotypes (*F*_1,19_ = 18.13, *q* < 0.001; *post-hoc*: between riverine and anadromous, *p* < 0.001, and riverine and potamodromous, *p* < 0.001) (Fig. 6B; Supplementary material: Table S21). This module had 12 enriched GO terms; genes were related to stimulus response, cell cycle, process and organisation (Supplementary material: Table S22, Fig. S2), amongst them *Fads2* on chromosome 4. Riverine trout had significantly lower module eigengene values than either migratory ecotype in another co-expression module, arbitrarily designated ‘blue’, with 1260 genes, including *Elovl2* on chromosome 2 and *Elovl5* on chromosome 18 (*F*_1,19_ = 8.68, *q* = 0.007; *post-hoc*: between riverine and anadromous, and riverine and potamodromous, both *p* = 0.005) (Fig. 6B; Supplementary material: Table S21). The ‘blue’ module showed 24 enriched GO terms, and most genes were related to metabolic and biosynthetic processes (Supplementary material: Table S23, Fig. S2). The ‘lightsteelblue’ module encompassed too few genes to perform a GO enrichment analysis. However, it showed lower module eigengene values in riverine trout than in either migratory type (*F*_1,19_ = 14.5, *q* = 0.001; *post-hoc*: between riverine and anadromous, and riverine and potamodromous, both *p* < 0.001), and it included a second *Fads2* locus on chromosome 4 (Fig. 6B; Supplementary material: Table S21).

**Fig. 6.**
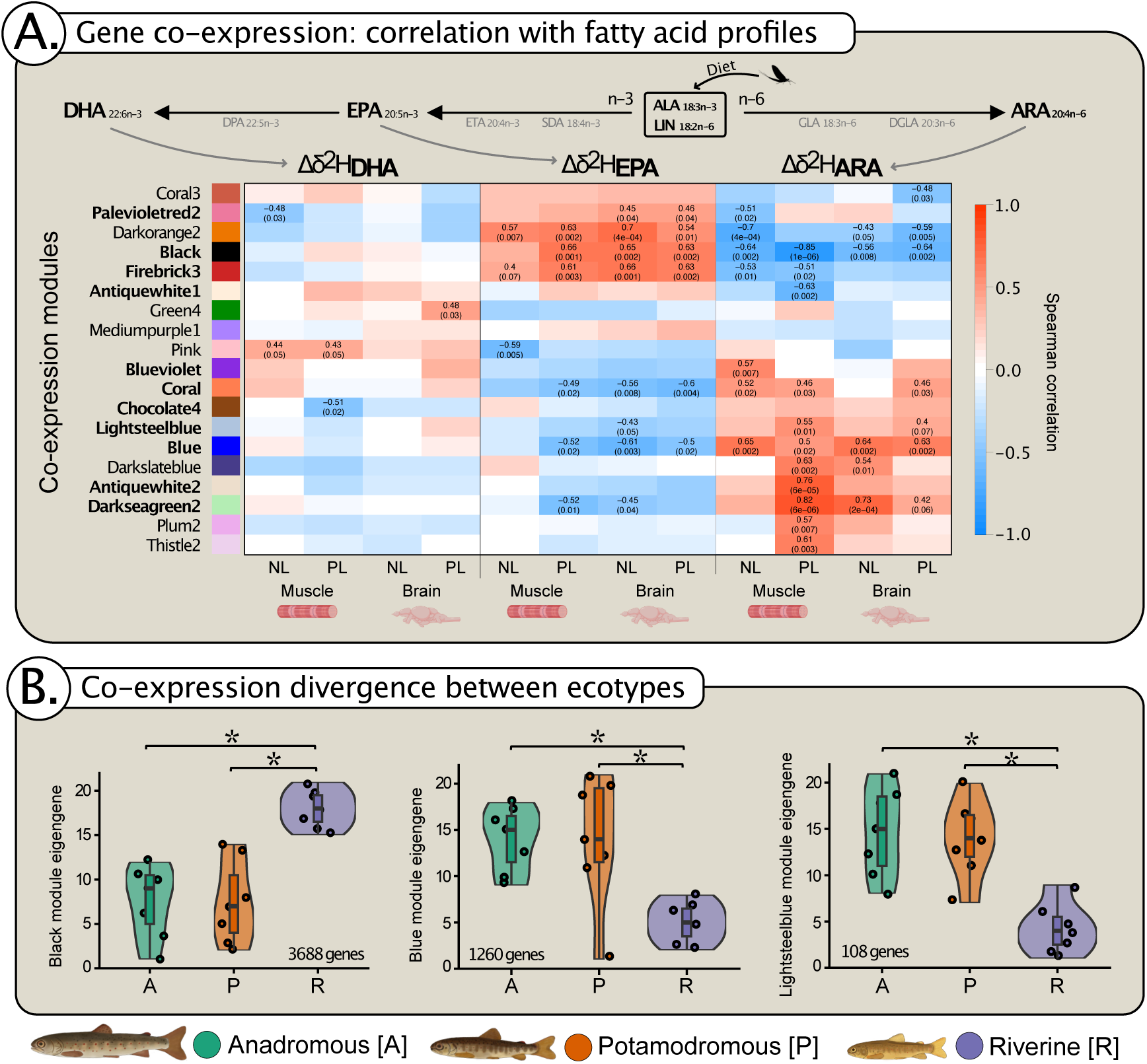
Gene co-expression modules for three brown trout ecotypes. **A.** Spearman’s rank correla,on coefficients with *p*-values in parentheses showing posi,ve (red) and nega,ve (blue) correla,ons of gene co-expression modules (given arbitrary colour names) with at least some LC-PUFA Δ*δ*^2^H profiles across neutral lipids (NL) and polar lipids (PL) in muscle and brain,ssue; modules in bold are also significantly associated with ecotype. **B.** Gene co-expression module eigengenes for modules associated with ecotype dis,nc,on and LC-PUFA biosynthesis, *and* containing *Fads2*, *Elovl5* and/or *Elovl2* genes. Asterisks (*) indicate significant differences between ecotypes.

### SNPs across all expressed genes

To evaluate potential genetic differentiation between ecotypes, we called SNPs across all (24,993) genes expressed in the liver. There was clear distinction between riverine and the migratory types (riverine *vs*. potamodromous mean *F_ST_* = 0.1041; riverine *vs.* anadromous mean *F_ST_* = 0.0997), but less between anadromous and potamodromous, with mean *F_ST_* effectively zero (Supplementary material: Figs. S3, S4).

### Candidate gene expression in common garden experiment

To infer whether differential regulation of *Fads2* and *Elovl5* genes truly represents responses to a dietary lack of n-3 LC-PUFA that helps to define migratory ecotype, we re-analysed mRNAseq data from an experimental population of Irish brown trout ecotypes that were reared on a common LC-PUFA-rich diet as a comparison dataset. In contrast to the wild trout of the Endrick Water, among the experimental trout of Wynne et al. (*29*), there was no significant difference in the expression of *Fads2* at any locus between ‘residents’ and ‘smolts’. Similarly, there was no significant difference between ecotypes in the expression of the *Elovl5* gene on chromosome 18. Only *Elovl5* on chromosome 10 was up-regulated in residents (LFC = 1.25 ± 0.44 SE, *p* = 0.013).

## Discussion

Analysing compound-specific stable hydrogen isotopes (*δ*_2_H) in fatty acids, coupled with lipid classes and gene expression analyses, revealed that juvenile brown trout compensate for deficiency in dietary n-3 LC-PUFA. Fish from all three ecotypes synthesised DHA by converting precursors, indicated by isotopic depletion of Δ*δ*^2^H_DHA_ values. Juvenile brown trout also routed DHA to polar lipids, where they are used as constituents of cell membranes. However, riverine residents and migrants displayed LC-PUFA synthesis and routing to different degrees, suggesting that ecotypes have different n-3 LC-PUFA requirements. The riverine ecotype appears to have a higher capacity to convert precursors to DHA, but also seems to have a lower DHA requirement, evidenced by lower DHA contents in polar and neutral lipids and tissue types (brain and muscle) than migrants, which could constitute excellent adaptations for life in headwaters with diets deprived of n-3 LC-PUFA. The importance of n-3 LC-PUFA in the divergence of life-history strategies have three main indicators, revealed by this study: 1) each ecotype was distinguished by its unique LC-PUFA composition and Δ*δ*_2_H values; 2) gene co-expression modules showed clear associations with both LC-PUFA biosynthesis and ecotype; and 3) genes directly involved in PUFA conversion (*Fads2* and *Elovl5*) were significantly up-regulated in riverine residents compared to migrant ecotypes.

Because vertebrates must obtain ALA or LIN from their diet (*21*, *22*), differences in the *δ*_2_H values suggest different dietary sources of these two essential PUFA. Although it is not possible to deduce the precise diets of each ecotype, the lower EPA and ARA contents of riverine trout compared to migrants are at least consistent with a higher proportion of food of terrestrial origin (*28*, *30*). Naturally, this is complicated by biosynthesis of these fatty acids (Supplementary material; Tables S14, S15, S24 and S25). Greater reliance on terrestrial input in riverine trout is supported by lower *δ*^2^H_ALA_ values but confounded by higher *δ*^2^H_LIN_ values (*31*).

The conclusion that diets in early life differed by trout ecotype must further be tempered with the acknowledgement that differing *δ*_2_H values may originate at the base of the food web (*32*). Because riverine trout were obtained from above waterfalls, it is likely, given the diverse *δ*^2^H_ALA_ and *δ*^2^H_LIN_ values, that there may be different sources of dietary ALA and LIN above and below the falls. Therefore, it may that the prey consumed basal food sources of different isotopic values that are cascaded up to trout. Furthermore, although there was no difference in the allocation of LIN to PL or NL between ecotypes, riverine trout allocated more ALA to muscle PL and, especially, muscle NL than migrant types did (Supplementary material: Tables S5–S7), the retention of which may skew *δ*^2^H_ALA_ values (*10*, *12*).

Across all ecotypes, DHA in trout was clearly more strongly distributed to cell membranes (PL; i.e., readily used) rather than to storage lipids (NL) in brain and muscle tissues, suggesting priority routing of available DHA to membrane buildup (*20*). However, such DHA routing did not seem to favour brain over muscle tissues. The use of DHA is not exclusively tied to neural development, and its significant presence in muscle tissue may be related to growth and function (*33*). Significantly depleted δ^2^H_DHA_ values revealed that of three ecotypes of trout synthesised DHA from precursors. Although riverine trout showed generally lower DHA contents, compared to either migratory type, these were not reflected in correspondingly lower Δ*δ*^2^H_DHA_ values. This indicates that all trout had obtained enough DHA for brain and muscle development via biosynthesis. Partial compensation for low dietary n-3 LC-PUFA supply via bioconversion of short-chain PUFA has previously been established experimentally in rats (*34*) and humans (*35*). Similarly, all brown trout ecotypes here appear to have compensated effectively for their dietary lack of DHA.

The more negative levels of Δ*δ*^2^H_ARA_ in riverine trout indicate higher ARA biosynthesis compared to either migratory type. Omega-6 PUFA, and ARA in particular, are important for inflammation, stress resistance, and osmoregulation (*36*). However, consideration of n-6 PUFA cannot be divorced from n-3 PUFA: both ALA and LIN substrates compete for the same enzymes during endogenous conversion to EPA or ARA (*37*). The bioconversion of n-3 PUFA may be generally higher than that of n-6 PUFA, as suggested by a ∼2.5:1 ratio in zebrafish (*Danio rerio*) (*38*). Thus, the abundance of biosynthesised ARA in riverine trout suggests even greater conversion of DHA and EPA if these PUFA are equally low in dietary supply.

This study has established that much of the differentiation in gene expression, in terms of individual candidate genes and networks of co-expressed genes, was clearly associated with traits of both migratory ecotype and LC-PUFA biosynthesis. The gene expression directly involved in LC-PUFA synthesis helps clarify LC-PUFA profiles among ecotypes, with the important caveats that the duration and stability of observed expression patterns remain unclear. Nonetheless, *Fads2* and *Elovl5* were both significantly up-regulated among riverine trout compared to either migrant type. We conclude that riverine trout biosynthesised appreciably more LC-PUFA than either migratory type. Otherwise, gene expression, generally, showed clear distinctions between riverine trout and both migratory ecotypes, in accordance with other studies (*29, 39*), but relatively little between anadromous and potamodromous.

The up-regulation of *Fads2* expression in the wild riverine residents compared to migratory trout was not replicated in the residents compared to smolts in the common-garden experiment by Wynne et al. (*29*). Although Wynne et al. (*29*) found differential expression between ecotypes in genes associated with metabolic pathways underlying energetic requirements, our re-analysis revealed no significant difference in *Fads2* expression. These experimental trout had all been raised on commercial pellets (*29, 40*), which typically contain marine-derived ingredients, including fish oil, rich in n-3 LC-PUFA (*41*), and likely negate the need for bioconversion. This suggests that the conversion of short-chain to long-chain PUFA seen in the wild riverine residents is a plastic response to dietary deficiency.

Although, in our study, the differentiation in gene expression was clear between riverine residents and migratory types, it remains unclear whether this is a driver or a result of the alternative migratory strategies (*29*). It has been suggested that alternative migratory strategies in salmonids may, in part, be the result of differential expression in early ontogeny of ‘master regulator’ genes (*42*), which causes divergent phenotypes and may precipitate such differences as observed here (*5*). It was also clear from SNP-calling that the ecotypes were genetically distinct, particularly the riverine resident from the migrant ecotypes. Capturing riverine trout from above a barrier ensured that they were truly resident, and not merely migrants in waiting, but our differentiation analyses suggest the barrier also provided a degree of reproductive isolation (*43*). It should be noted that the extent to which the differentiation in migratory and fatty acid traits is due to population effects still requires further investigation.

The size difference between ecotypes is consistent with the hypothesis that increased LC-PUFA biosynthesis among riverine trout diverts energy from somatic growth (*11*). Because all specimens were the same age, contrasting growth rates can reasonably be inferred from fork lengths, which differed markedly among ecotypes, although size at hatching remains, naturally, unknown, as do parental effects such as gamete provisioning (*44*). Previous studies reported variable body size among juvenile brown trout between ecotypes: some found the fastest growing trout tended to remain as river residents (*45, 46*), while others argued the opposite to be the case (*5*, *47*). Despite their smaller size, riverine trout showed much higher total muscle lipid content than anadromous or potamodromous. Future migrants, but not riverine residents, may compensate for the slowing of growth due to winter starvation by allocating resources to protein metabolism at the expense of lipids (*48*). Earlier studies demonstrated a transcriptomic link with depleted lipid stores associated with the smolting process in salmonids (*49, 50*). It seems likely that responses to winter starvation, individual variation in physiology and changes related to smolting, combined with diet differences, account for discrepancies in total lipids between these riverine and migratory ecotypes of brown trout (*2*). We conclude that riverine residents sacrifice somatic growth to invest more energy into lipid production and maintenance, including LC-PUFA biosynthesis, than migrants do.

Migration confers higher fitness benefits on female than on male salmonids (*51, 52*) with differences between sexes in juvenile physical characteristics and gene expression (*53, 54*). Females may well have a greater propensity to migrate than males to take advantage of the future larger body size and greater fecundity associated with migration to more nutrient-rich habitats (*47*). However, sex-specific effects on any of the variables we tested were minor and likely due to the age of the trout (2+ cohort), before sexual maturity (*29*, *55*).

The similarity in fatty acid profiles and gene expression between anadromous and potamodromous ecotypes suggests that migration *vs* residency is a more fundamental life-history distinction than anadromy *vs* non-anadromy. It remains unclear why migratory trout down-regulated expression of genes associated with PUFA conversion enzymes, yet they had more n-3 LC-PUFA in their brains than did riverine residents. It is possible that migratory fish increase their conversion capacity through early ontogeny to reach a specific n-3 LC-PUFA level required for successful migration, or retain these nutrients from the egg stage. The latter argument would require compound-specific stable isotope analysis of PUFA from eggs to track maternal n-3 LC-PUFA carryover to fish organs. Alternatively, migratory trout may selectively retain dietary n-3 PUFA more efficiently that residents. Regardless, it appears that low n-3 LC-PUFA does not drive individuals to migrate, but rather a lack of capacity to maintain levels of n-3 LC-PUFA synthesis sufficient to spend their whole lives in n-3 LC-PUFA-deprived headwaters.

This is the first study that associates life-history ecotype with LC-PUFA bioconversion, underpinned by differences in gene expression at the point at which first outward migration occurs. Future studies are warranted to determine the extent to which the results presented here are replicable across various systems and lineages of brown trout. Consideration should also be given to different age classes, especially earlier stages of ontogeny, to assess whether and how the suggested associations are causes or consequences of life-history divergence, and effects of dietary and biosynthesised n-3 LC-PUFA on brain development, cognitive behaviour and physiology between brown trout ecotypes (*11*). Our results suggest that capacity to compensate for lack of dietary DHA in nursery streams through biosynthesis and routing, needed for brain nutrition and cognitive function, is crucial to the life-history decisions of juvenile brown trout.

## Methods

### Experimental design

We collected three migratory ecotypes of a cohort of wild juvenile brown trout from a single nursery stream, at their first outward migration, with the objectives: a) to investigate whether migratory ecotypes can be characterised by total lipids, dietary fatty acids, and contents of LC-PUFA in brain and muscle lipids; b) to determine whether there is an association of ecotype and LC-PUFA biosynthesis, and whether this is reflected in divergence in somatic growth; and c) to evaluate whether LC-PUFA biosynthesis is in response to dietary deficiency.

We measured fork lengths of anadromous, potamodromous and riverine resident trout of the same age to assess and compare growth rates. We extracted lipids from brain and muscle tissue, and isolated individual n-3 and n-6 PUFA from polar and neutral lipids to compare lipid and fatty acid contents between ecotypes. We used compound-specific SIA (*δ*^2^H) of individual PUFA along the n-3 and n-6 conversion pathways to compare signatures of biosynthesis. We sequenced mRNA from liver tissue to perform a gene expression analysis and construct modules of co-expressed genes. We focussed on genes known to be involved in fatty acid conversion pathways to uncover differences between ecotypes. We also called SNPs to determine genetic differentiation between ecotypes. Finally, we compared our data to the RNAseq data from another study, which used a single LC-PUFA enriched common-garden experimental diet to infer whether LC-PUFA biosynthesis in the wild trout was likely to be in response to dietary lack.

A role for sex in routing and synthesis of n-3 LC-PUFA in some fish species (*e.g.* Eurasian perch *Perca fluviatilis*) (*56*). Although sex differences in brown trout are not expected before maturity (*55*), we considered the potential impact of sex in all of our analyses.

### Sample collection

Samples (*n* = 85) of three migratory ecotypes of brown trout were collected from the Endrick Water, a fourth-order stream in the Leven (Strathclyde) hydrological area (HA85), Scotland, UK, which flows into Loch Lomond and, thence, via the River Leven, to the Clyde Estuary and the sea (Fig. 1). All trout were captured at the time of first outward migration, March to May 2021. Anadromous (females *n* = 10, males *n* = 21) and potamodromous (females *n* = 15, males *n* = 13) types were caught with a rotary screw trap, placed near the point at which no further suitable salmonid habitat existed downstream, and identified as smolts (including pre-smolts) or parr using the morphological criteria described by Tanguy et al. (*57*). To confirm that putative potamodromous parr were, indeed, migrating, specimens were marked with visible implant elastomer (Northwest Marine Technology Inc.), transported *ca.* 5 km upstream, released and recaptured at the trap within 24 hours. Riverine resident trout (females *n* = 8, males *n* = 18) were electro-fished above an ‘impassable’ barrier (waterfalls) to ensure they were unable, or at least unlikely, to migrate into lentic waters.

Specimens were killed by benzocaine overdose, fork-length measured (*i.e.* from tip of snout to fork in tail), and digitally photographed on the left side. Morphological differences between ecotypes were confirmed with comparisons of fork length and geometric morphometric analyses (Supplementary material: text; Figs. S5, S6; Tables S26, S27). All trout were aged by scale examination, and only those judged to be in the 2+ cohort (*i.e.* third spring after hatch) were retained for investigation. Adipose fin clips were used as a source of nuclear DNA with which to genotype for sex following a modified duplex PCR protocol (Supplementary material: text; Tables S28–S30). Seven specimens of each ecotype and roughly equal numbers of each sex were selected at random for further analyses: whole brains and samples of dorsal muscle tissue were flash frozen, then freeze-dried for fatty acid analyses, while livers were preserved in RNA*later* (Sigma-Aldrich) for RNAseq analyses.

### Fatty acid analysis

The protocol described by Pilecky et al. (*58*) was followed for lipid extraction, esterification, and separation into polar (PL) and neutral lipids (NL) from freeze-dried samples of brain and muscle tissue from the same subset of 21 specimens described above. Gas chromatography (TRACE GC 1310, Thermo) was used to separate and quantify fatty acid methyl esters (FAME) of PL and NL using external standards. FAME were reported as mg g^−1^ sample dry weight (*58*). Fatty acid-specific stable isotopes were measured using gas-chromatography (see above) coupled with an isotope ratio mass spectrometry (DELTA V Advantage, ThermoFisher Scientific) via CONFLO IV (Thermo) and compared with certified ME-C20:0 stable isotope reference material: USGS70: *δ*^2^H = −183.9 ‰, USGS71: *δ*^2^H = −4.9 ‰ and USGS72: *δ*^2^H = +348.3 ‰. Samples were corrected for methylation as described by Pilecky et al. (*58*).

ALA and LIN stand at the base of the n-3 and n-6 bioconversion pathways, respectively. Therefore, group differences in dietary nutrition were determined by testing with ANOVA the effects of ecotype and sex on the *δ*^2^H of ALA and LIN for each lipid class and tissue type (*i.e.* PL and NL from brain and muscle); Tukey’s HSD followed *post hoc*. Because exact diets of individual wild trout were unknown, correction to *δ*^2^H signatures in accordance with differences in diet was accomplished for each LC-PUFA with reference to a fatty acid preceding it in the appropriate synthesis pathway:

1. Δ*δ*^2^H_EPA_ = sample *δ*^2^H_EPA_ – mean(specimen *δ*^2^H_ALA_),
2. Δ*δ*^2^H_DHA_ = sample *δ*^2^H_DHA_ – mean(specimen *δ*^2^H_EPA_),
3. Δ*δ*^2^H_ARA_ = sample *δ*^2^H_ARA_ – mean(specimen *δ*^2^H_LIN_),

in which specimen refers, in this context, to each lipid class and tissue type from one individual, and the LC-PUFA sample in question is from the same individual. Bioconversion of shorter chain PUFA to LC-PUFA, instead of a dietary source of LC-PUFA, is indicated by depletion of Δ*δ*^2^H in the applicable FAME.

### RNA sequencing and gene expression analyses

High-quality RNA was extracted from the liver tissue of the subset specimens, using an Invitrogen™ PureLink™ RNA Mini Kit (ThermoFisher Scientific), following the manufacturer’s instructions, except for an additional homogenisation that used FastPrep-24 (MP Biomedicals) before isolation. RNA quantity was assessed using the Qubit 2.0 fluorometer (ThermoFisher Scientific) and quality using 2200 Tapestation (Agilent, Santa Clara, CA). Ratios of A260/280 were between 2.1 and 2.2, while RNA Integrity Numbers were 8.8 or higher.

Individually barcoded poly(A) mRNAseq standard library preparation was done from 1 µq of high-quality RNA either in-house (11 samples; using the NEBNext® Ultra™ II Directional RNA Library Prep Kit for Illumina® with the NEBNext® Poly(A) mRNA Magnetic Isolation Module) or at Novogene UK (10 samples). Libraries were sequenced to 30M 150bp paired-end reads per sample at Novogene UK (Cambridge) on two runs using Illumina NovaSeq 6000 (Supplementary material: Table S31).

Untrimmed reads (maximum read length = 150 bp) were aligned to the brown trout reference genome (fSalTru1.1, INSC Assembly GCA_9010011651.1) (*59*) and annotated for novel splice junctions using *STAR* v. 2.7.10b (*60*) in a two-step mapping approach. Duplicates were marked with the *Picard* tool (https://broadinstitute.github.io/picard). Read counts were generated with *HTSeq* v. 2.0 (*61*) and processed using *DESeq2* v. 3.17 (*62*); transcripts with less than 10 reads in at least 7 individuals were excluded from the analysis.

Gene co-expression networks were constructed, based on *rlog*-transformed read counts using *WGCNA* v. 1.12-1 (*63, 64*). Modules were determined with a dynamic tree algorithm, with a minimum size of 30 genes, and were named after arbitrary colours. Similar modules were merged based on eigengene distance threshold of 0.25.

For each co-expression module associated with ecotype and LC-PUFA conversion, of which candidate *Fads* and *Elovl* genes were part, gene ontology (GO) enrichment analyses were conducted with *g:Profiler* v. e110_eg57_p18_4b54a898 (*65*) and summarized with the *R* package, *simplifyEnrichment*, using the ‘binary cut’ clustering algorithm (*66*). The GO-annotation used was from *Ensembl* v. 110 (*67*), included with *g:Profiler*, and the background dataset for the GO enrichment consisted of all *expressed* genes, rather than all genes in the genome.

### SNP calling

SNPs were called from trimmed reads with the following approach: raw reads were processed with *Trimmomatic* v. 0.39 (*68*), leading and trailing bases with a Phred score below 20 were discarded, and a sliding 4-bp window approach was employed to trim reads where Phred scores dropped below 20. SNP calling was conducted using *FreeBayes* v.1.3.7 (*69*), using a minimum coverage of 3. The resulting dataset was filtered using *BCFtools* (*70, 71*), retaining biallelic SNPs with a minimum genotype depth of 5, genotype quality of 20, less than 20% missing data, and minor allele frequency above 10%.

### Comparison data

Although their study did not investigate LC-PUFA bioconversion, the differential gene expression data published by Wynne et al. (*29*) provided a useful point of comparison with the present study. Their RNAseq data derived from 24 brown trout liver samples were downloaded from the NCBI Sequence Read Archive (BioProject ID: PRJNA670837) using *fastq-dump* in *SRA Toolkit* (*72*) with the --split-3 command to separate reads into forward and reverse. We then conducted alignment of untrimmed reads (max. 150 bp) to the reference genome and splice junctions, read-count generation and processing.

Wynne et al. (*29*) raised brown trout in captivity from egg to age 2+. Trout were the offspring of wild-caught parents (4 females and 5 males) from a single population in the Republic of Ireland, known for high rates of anadromy. They formed 8 full-sibling and 4 half-sibling families. All fish were reared on the same LC-PUFA-rich commercial pellet diet, fed *ad libitum* (*22*, *40*). Over 22 months, fish were assessed morphologically for smolting and tested for salt-water tolerance, and classed into three groups with females and males in roughly equal proportions: putative smolts exposed to salt water (*n* = 7), putative residents exposed to salt water (*n* = 9), and putative residents not exposed to salt water (*n* = 9).

### Statistical analysis

Unless otherwise stated, *R* v.4.2.2. (*73*) was used for all statistical analyses. We modelled and tested effects of ecotype and sex on fork length with ANOVA, followed by Tukey’s HSD *post hoc*.

We used MANOVA to test the effects of ecotype and sex on total (polar and neutral) lipids in brain and muscle tissue. ANOVA followed by Tukey’s HSD *post hoc* were then used to examine total lipids in tissues individually by ecotype. We fitted linear mixed-effects models with the *R* package *lme4* (*74*), which employs Satterthwaite’s method of calculating degrees of freedom, to examine differences in relative content of each FAME between lipid classes and tissue types (*i.e.* the effect of lipid class and tissue type on FAME percentage) across all specimens, with individual specimen treated as a random effect. We tested pairwise differences between lipid classes and tissue types *post hoc* with Tukey’s method. We tested ecotype and sex effects on the relative content of individual LC-PUFA with ANOVA for each lipid class and tissue type followed by Tukey’s HSD *post hoc*. We used the same procedure to test effects on Δ*δ*_2_H. Finally, to inform probability of ecotype based solely on LC-PUFA signatures, we fitted multinomial logistic regressions, using the *R* package *nnet* (*75*), in various combinations:

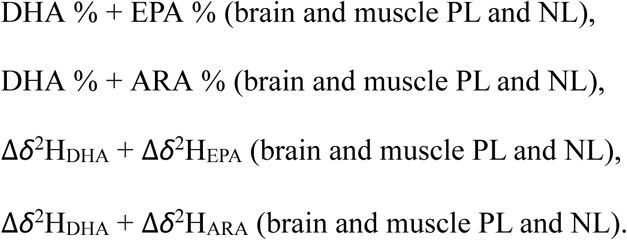

We performed principal components analysis (PCA) on *rlog*-transformed read counts from RNAseq using the *R* package, *pcaMethods* (*76*), with the settings: scaling = “none”, center = TRUE. We identified genes differentially expressed by ecotype and sex using pairwise analyses (Anadromous : Riverine, Anadromous : Potamodromous, Potamodromous : Riverine, and Female : Male) and tested them with Wald tests in *DESeq2* v. 3.17 (*62*), selecting genes with *z*-transformed loadings above 2 or below −2, corresponding to a *p*-value of 0.05.

We tested relationships between gene co-expression modules and ecotype and sex with ANOVA, and discovered those between modules and Δ*δ*_2_H of EPA, DHA and ARA using Spearman’s rank correlation coefficient.

We performed PCA on linkage-disequilibrium-pruned SNPs in the *R* package, *SNPRelate* (*77*), only retaining SNPs with linkage disequilibrium (r^2^) below 0.2. We calculated weighted *F_ST_* genome-wide and on a SNP-by-SNP basis using *VCFtools* (*78*) and visualised these with the *R* package, *CMplot* (*79*).

For the comparison RNAseq dataset of Wynne et al. (*29*), we performed PCA on *rlog*- transformed read counts and identified by pairwise analysis and testing, as above, genes differentially expressed between smolts and residents, with exposure to salt water treated as a covariate.

## Supporting information

Supplementary results

TableS20-Gene_expression_results

TableS21-Co-expression-results

## Acknowledgements

The authors are grateful to the teams at the Scottish Centre for Ecology and the Natural Environment (SCENE) and the Loch Lomond Fisheries Trust for assistance in the field and to Maria Capstick and Samuel-Karl Kämmer for laboratory assistance. All work with live specimens was conducted under UK Home Office Licence No. 70/8794.

## Funding

Natural Environment Research Council Independent Research Fellowship NR/W008963/1(AJ)

Leverhulme Trust Early Career Fellowship ECF-2020-509 (AJ)

Austrian Science Fund (FWF) [10.55776/P35515] (LZ)

Fisheries Society of the British Isles PhD studentship (JPK, hosted by CEA, KRE)

## Author contributions

Conceptualisation: JPK, LZ, CEA

Data curation: JPK

Formal analysis: JPK, MP

Funding acquisition: AJ, LZ, CEA

Investigation: JPK

Methodology: AJ, LZ, MP, MJK

Project administration: JPK, HMH

Resources: AJ, MJK, KRE, CEA

Supervision: LZ, KRE, CEA

Writing – original draft: JPK

Writing – review & editing: JPK, AJ, LZ, MP, HMH, MJK, KRE, CEA

## Data and materials availability

All data are available in the main text or the supplementary materials, except for additional data and code deposited at https://figshare.com/s/893a81181cc5c8f58c07 (private link to be made public upon publication), and cDNA sequences, which will be uploaded to NCBI GenBank upon publication. Access to data from Wynne et al., 2021 is indicated in the reference provided (*29*).

