## Supplementary results for "A critical role for brain nutrition in the life-history decisions of a partially migratory fish"

### Supplementary materials

### Supplementary text

#### *Geometric morphometric analysis*

Immediately upon death by benzocaine overdose, all trout were digitally photographed on the left side. Images were converted to thin plate spline files with tpsUtil (*1*). One researcher then scale-calibrated and landmarked 13 homologous points using tpsDig2 (*2*) (Fig. S5). A generalised Procrustes analysis with a 1,000-round randomised residual permutation procedure was performed in the *R*-package, 'Geomorph v.3.1.2' (*3*), and the effects of centroid size and ecotype on body shape was tested using Procrustes ANOVA (Table S26). Possible effects of allometry were accounted for by using the residuals of a regression of Procrustes coordinates on centroid size; the effectiveness of the account was confirmed with MANOVA testing the effect of fork length on the residuals and finding no significant effect ($Pillai = 0.376$, $F_{1,83} = 1.51$, $p = 0.102$). Least-squares means distance comparisons, derived from the permutation procedure above with 95% confidence, were performed with the 'pairwise' function in 'RRPP' (*4*); all pairwise comparisons of ecotypes were highly significant (Table S27). Each ecotype had its own characteristic morphology (Fig. S6).

#### *Genotyping for sex*

Following the manufacturer's instructions, a NucleoSpin™ Tissue kit (Macherey-Nagl) was used to extract genomic DNA from adipose fin clips. Quality was controlled with spectrophotometry (NanoDrop, ThermoFisher Scientific), and quantification was conducted fluorometrically (Qubit 2.0, ThermoFisher Scientific), before dilution to 20 ng $\mu L^{-1}$. Duplex PCR amplified the male-chromosome gene, *sdY*, with *18S* as a positive amplification control (Tables S3–S5), and agarose gel (2 %) electrophoresis was used to visualise resulting products (*4–5*).

Supplementary Figures

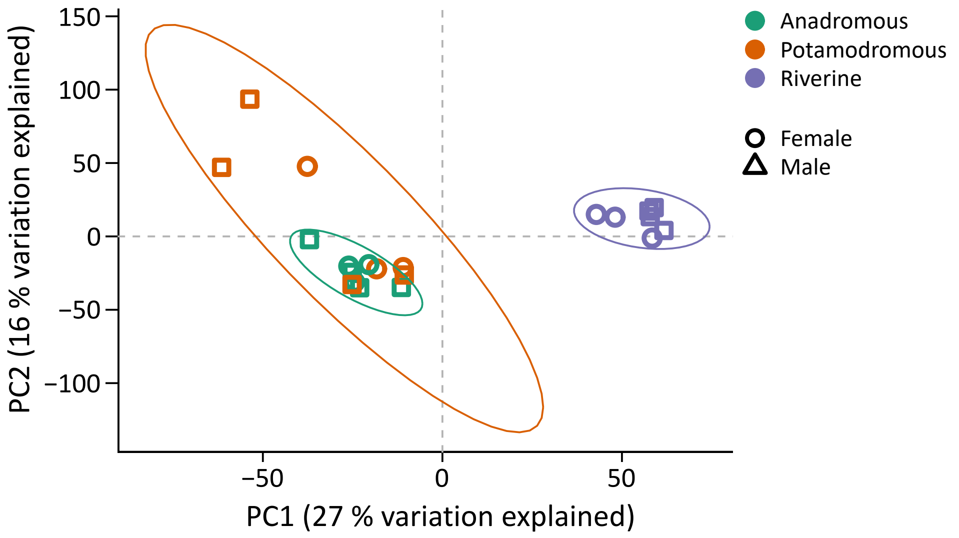


**Fig. S1.** PCA of *rlog*-transformed read counts indicating differences in gene expression patterns in liver tissue between three ecotypes of brown trout.

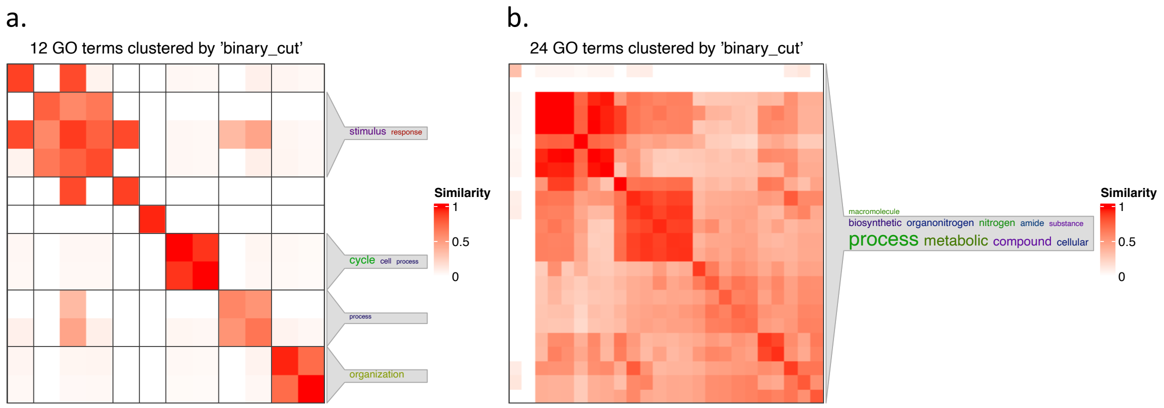


**Fig. S2.** Gene ontology terms cluster by 'binary_cut' for a. 'black' and b. 'blue' gene co-expression modules in brown trout liver tissue.

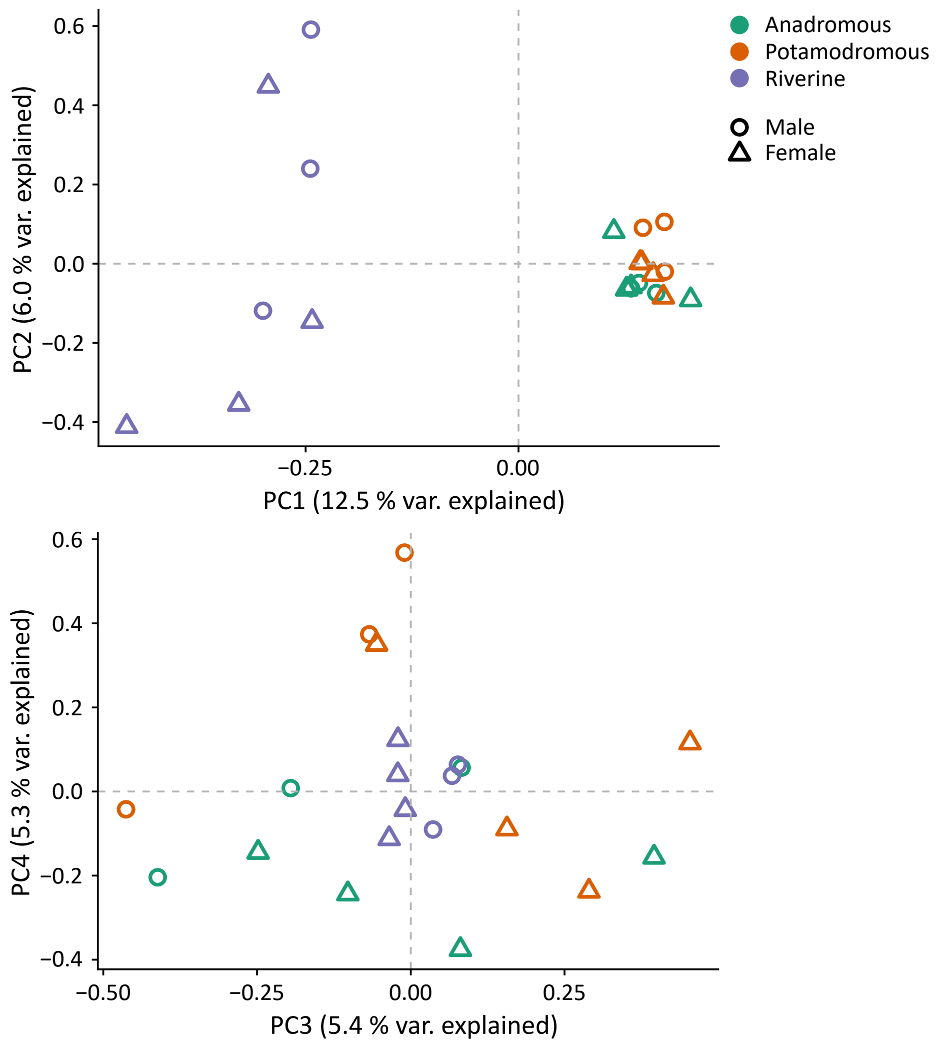


**Fig. S3.** PCA plots of SNPs revealed by RNAseq for three ecotypes of brown trout.

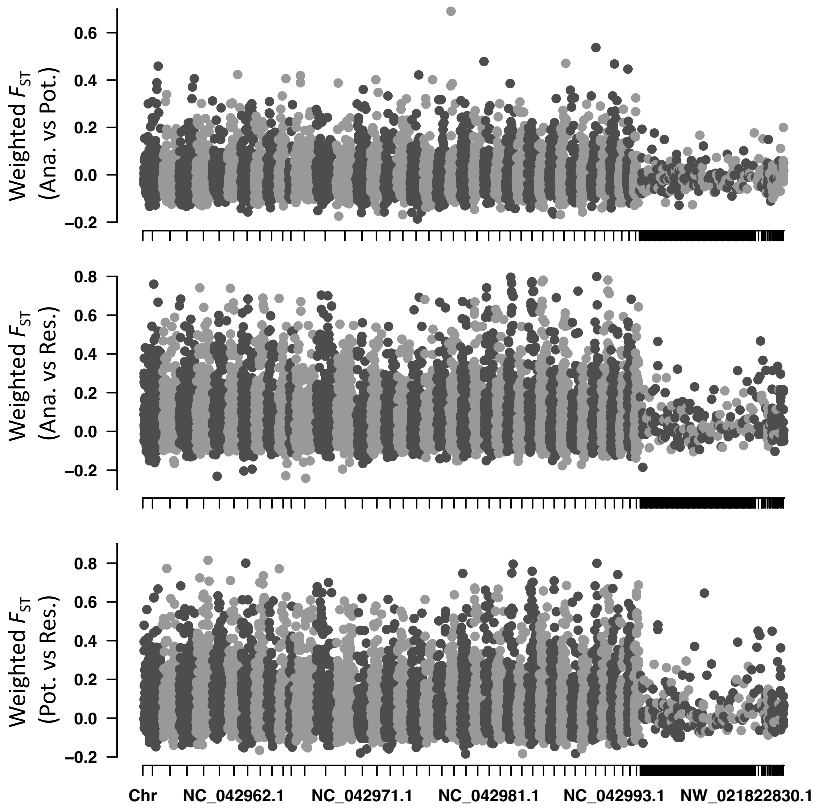


**Fig. S4.** Manhattan plots of pairwise weighted $F_{ST}$ of SNPs in RNAseq data from three ecotypes of brown trout.

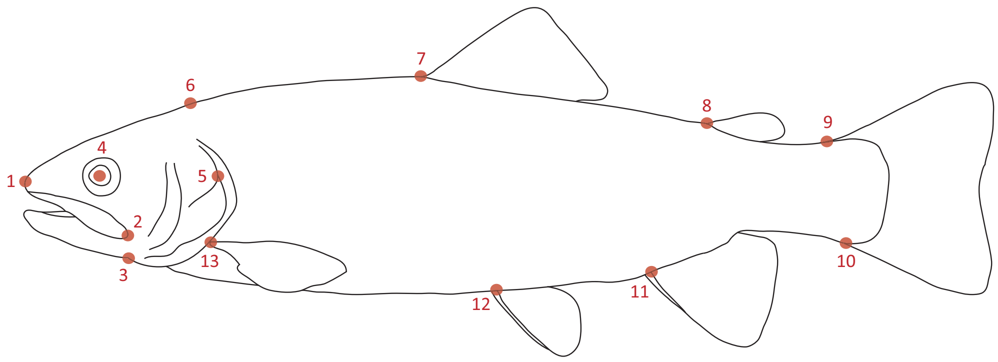


**Fig. S5.** Landmarks used for geometric morphometric analysis of brown trout shape: 1. tip of snout; 2. posterior of maxilla; 3. joint of lower jaw; 4. centre of eye; 5. anterior intersection of opercula and subopercule; 6. superior posterior of cranium; 7. anterior insertion of dorsal fin; 8. anterior insertion of adipose fin; 9. dorsal junction of caudal fin; 10. ventral junction of caudal fin; 11. anterior insertion of anal fin; 12. anterior insertion of pelvic fin; 13. superior insertion of pectoral fin.

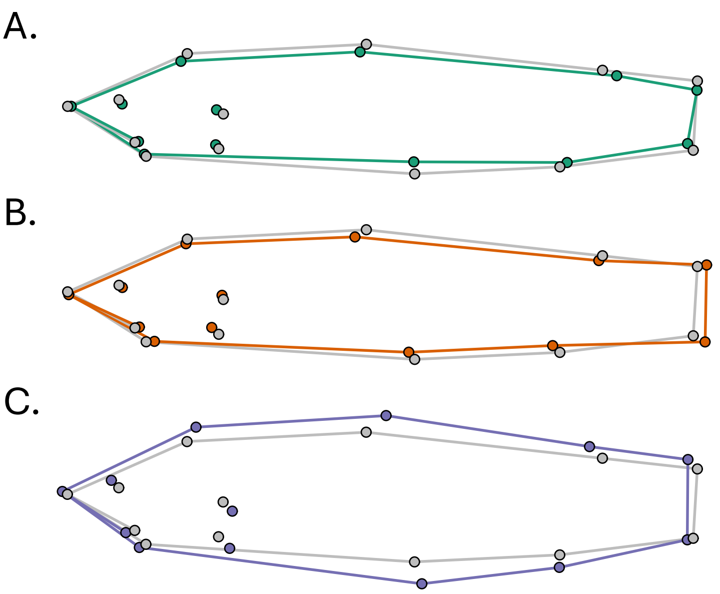


**Fig. S6.** Consensus shapes of A. anadromous, B. potamodromous, and C. riverine brown trout from Endrick Water, Scotland, overlaid on the consensus shape of all (grey). For clarity, shape differences are shown with 3x exaggeration.

Supplementary Tables

**Table S1.** ANOVA results for effects of ecotype and sex on $\delta^2H_{ALA}$ in two lipid classes from two tissue types. PL = polar lipids, NL = neutral lipids

| | | **df** | **Sum Sq.** | **Mean Sq.** | ***F*** | **Pr(>*F*)** |
|---|---|---|---|---|---|---|
| Brain PL | Ecotype | 2 | 7255 | 3627 | 7.05 | 0.021 |
| | Sex | 1 | 3 | 3 | 0.01 | 0.942 |
| | Residuals | 7 | 99.0 | 514 | | |
| Brain NL | Ecotype | 2 | 1468 | 734 | 1.07 | 0.446 |
| | Sex | 1 | 683 | 683 | 1.00 | 0.392 |
| | Residuals | 3 | 2058 | 686 | | |
| Muscle PL | Ecotype | 2 | 124 | 62 | 0.26 | 0.772 |
| | Sex | 1 | 9 | 9 | 0.04 | 0.850 |
| | Residuals | 14 | 3290 | 235 | | |
| Muscle NL | Ecotype | 2 | 3380 | 1690 | 4.70 | 0.025 |
| | Sex | 1 | 79 | 79 | 0.22 | 0.646 |
| | Residuals | 16 | 5757 | 360 | | |

**Table S2.** Pairwise comparisons from Tukey's HSD *post hoc* tests of the effect of ecotype on $\delta^2H_{ALA}$ in brain and muscle lipids. PL = polar lipids, NL = neutral lipids

| | **Comparison** | **Difference** | **Lower** | **Upper** | ***p* adj** |
|---|---|---|---|---|---|
| Brain PL | Pot : Ana | -27.095 | -71.900 | 17.710 | 0.244 |
| | Riv : Ana | -73.675 | -131.518 | -15.832 | 0.017 |
| | Riv : Pot | -46.580 | -102.461 | -9.301 | 0.098 |
| Muscle NL | Pot : Ana | -7.495 | -34.725 | 19.735 | 0.761 |
| | Riv : Ana | -29.971 | -56.133 | -3.810 | 0.024 |
| | Riv : Pot | -22.476 | -49.706 | 4.754 | 0.115 |

**Table S3.** ANOVA results for effects of ecotype and sex on $\delta^2H_{LIN}$ in two lipid classes from two tissue types. PL = polar lipids, NL = neutral lipids

| | | **df** | **Sum Sq.** | **Mean Sq.** | ***F*** | **Pr(>*F*)** |
|---|---|---|---|---|---|---|
| Brain PL | Ecotype | 2 | 3929 | 1964 | 14.19 | 0.003 |
| | Sex | 1 | 68 | 68 | 0.49 | 0.506 |
| | Residuals | 7 | 969 | 139 | | |
| Brain NL | Ecotype | 2 | 615 | 308 | 1.14 | 0.380 |
| | Sex | 1 | 240 | 240 | 0.89 | 0.382 |
| | Residuals | 6 | 1616 | 269 | | |
| Muscle PL | Ecotype | 2 | 14696 | 7348 | 16.29 | < 0.001 |
| | Sex | 1 | 69 | 69 | 0.15 | 0.701 |
| | Residuals | 15 | 6767 | 451 | | |
| Muscle NL | Ecotype | 2 | 8110 | 4055 | 21.10 | < 0.001 |
| | Sex | 1 | 128 | 128 | 0.67 | 0.427 |
| | Residuals | 15 | 2882 | 192 | | |

**Table S4.** Pairwise comparisons from Tukey's HSD *post hoc* tests of the effect of ecotype on $\delta^2H_{LIN}$ in brain and muscle lipids. PL = polar lipids, NL = neutral lipids

| | Comparison | Difference | Lower | Upper | $p_{adj}$ |
|---|---|---|---|---|---|
| Brain PL | Pot : Ana | -1.813 | -27.122 | 23.496 | 0.976 |
| | Riv : Ana | 41.267 | 12.970 | 69.563 | 0.009 |
| | Riv : Pot | 43.080 | 17.771 | 68.389 | 0.004 |
| Muscle PL | Pot : Ana | 38.577 | 6.273 | 70.882 | 0.019 |
| | Riv : Ana | 64.471 | 34.981 | 93.961 | < 0.001 |
| | Riv : Pot | 25.894 | -6.410 | 58.199 | 0.127 |
| Muscle NL | Pot : Ana | 2.93 | -18.15 | 24.01 | 0.931 |
| | Riv : Ana | 43.99 | 24.74 | 63.23 | < 0.001 |
| | Riv : Pot | 41.05 | 19.97 | 62.14 | < 0.001 |

**Table S5.** ANOVA results for effects of ecotype and sex on ALA (18:3n-3) content (%) in two lipid classes from two tissue types. PL = polar lipids, NL = neutral lipids

| | | **df** | **Sum Sq.** | **Mean Sq.** | ***F*** | **Pr(>*F*)** |
|---|---|---|---|---|---|---|
| Brain PL | Ecotype | 2 | 0.24 | 0.12 | 1.84 | 0.189 |
| | Sex | 1 | 0.19 | 0.19 | 2.94 | 0.105 |
| | Residuals | 17 | 1.10 | 0.06 | | |
| Brain NL | Ecotype | 2 | 1.11 | 0.56 | 0.58 | 0.572 |
| | Sex | 1 | 2.70 | 2.70 | 2.80 | 0.113 |
| | Residuals | 17 | 16.39 | 0.96 | | |
| Muscle PL | Ecotype | 2 | 2.84 | 1.42 | 3.81 | 0.043 |
| | Sex | 1 | 0.08 | 0.08 | 0.21 | 0.653 |
| | Residuals | 17 | 6.33 | 0.37 | | |
| Muscle NL | Ecotype | 2 | 208.02 | 104.01 | 36.65 | < 0.001 |
| | Sex | 1 | 0.05 | 0.05 | 0.02 | 0.894 |
| | Residuals | 17 | 48.25 | 2.84 | | |

**Table S6.** Pairwise comparisons from Tukey's HSD *post hoc* tests of the effect of ecotype on ALA (20:3n-3) content (%) in muscle lipids. PL = polar lipids, NL = neutral lipids

| | Comparison | Difference | Lower | Upper | $p_{adj}$ |
|---|---|---|---|---|---|
| Muscle PL | Pot : Ana | -0.478 | -1.315 | 0.359 | 0.331 |
| | Riv : Ana | 0.421 | -0.416 | 1.258 | 0.419 |
| | Riv : Pot | 0.899 | 0.062 | 1.736 | 0.034 |
| Muscle NL | Pot : Ana | -0.569 | -2.289 | 1.741 | 0.805 |
| | Riv : Ana | 6.374 | 4.064 | 8.684 | < 0.001 |
| | Riv : Pot | 6.943 | 4.633 | 9.253 | < 0.001 |

**Table S7.** ANOVA results for effects of ecotype and sex on LIN (18:2n-6) content (%) in two lipid classes from two tissue types. PL = polar lipids, NL = neutral lipids

| | | **df** | **Sum Sq.** | **Mean Sq.** | ***F*** | **Pr(>*F*)** |
|---|---|---|---|---|---|---|
| Brain PL | Ecotype | 2 | 0.83 | 0.42 | 1.16 | 0.336 |
| | Sex | 1 | 0.44 | 0.44 | 1.22 | 0.285 |
| | Residuals | 17 | 6.07 | 0.36 | | |
| Brain NL | Ecotype | 2 | 10.87 | 5.43 | 1.30 | 0.298 |
| | Sex | 1 | 9.44 | 9.44 | 2.26 | 0.151 |
| | Residuals | 17 | 71.05 | 4.18 | | |
| Muscle PL | Ecotype | 2 | 0.45 | 0.22 | 0.49 | 0.623 |
| | Sex | 1 | 0.92 | 0.92 | 1.99 | 0.176 |
| | Residuals | 17 | 7.85 | 0.64 | | |
| Muscle NL | Ecotype | 2 | 14.74 | 7.37 | 2.11 | 0.152 |
| | Sex | 1 | 1.88 | 1.88 | 0.54 | 0.474 |
| | Residuals | 17 | 59.43 | 3.50 | | |

**Table S8.** ANOVA results for effects of ecotype and sex on EPA (20:5n-3) content (%) in two lipid classes from two tissue types. PL = polar lipids, NL = neutral lipids

| | | **df** | **Sum Sq.** | **Mean Sq.** | ***F*** | **Pr(>*F*)** |
|---|---|---|---|---|---|---|
| Brain PL | Ecotype | 2 | 21.5 | 10.8 | 25.66 | < 0.001 |
| | Sex | 1 | < 0.1 | < 0.1 | 0.01 | 0.924 |
| | Residuals | 17 | 32.2 | 1.9 | | |
| Brain NL | Ecotype | 2 | 85.8 | 42.9 | 22.64 | < 0.001 |
| | Sex | 1 | 0.2 | 0.2 | 0.10 | 0.751 |
| | Residuals | 17 | 32.2 | 1.9 | | |
| Muscle PL | Ecotype | 2 | 8.4 | 4.2 | 1.37 | 0.282 |
| | Sex | 1 | 1.7 | 1.7 | 0.55 | 0.467 |
| | Residuals | 17 | 52.2 | 3.1 | | |
| Muscle NL | Ecotype | 2 | 31.8 | 15.9 | 1.80 | 0.195 |
| | Sex | 1 | 3.1 | 3.1 | 0.36 | 0.556 |
| | Residuals | 17 | 150.2 | 8.8 | | |

**Table S9.** Pairwise comparisons from Tukey's HSD *post hoc* tests of the effect of ecotype on EPA (20:5n-3) content (%) in brain lipids. PL = polar lipids, NL = neutral lipids

| | **Comparison** | **Difference** | **Lower** | **Upper** | ***p* adj** |
|---|---|---|---|---|---|
| Brain PL | Pot : Ana | -0.723 | -1.611 | 0.165 | 0.122 |
| | Riv : Ana | -2.417 | -3.305 | -1.528 | < 0.001 |
| | Riv : Pot | -1.694 | -2.582 | -0.805 | < 0.001 |
| Brain NL | Pot : Ana | -2.467 | -4.355 | -0.579 | 0.010 |
| | Riv : Ana | -4.952 | -6.829 | -3.064 | < 0.001 |
| | Riv : Pot | -2.485 | -4.372 | -0.597 | 0.010 |

**Table S10.** ANOVA results for effects of ecotype and sex on ARA (20:4n-6) content (%) in two lipid classes from two tissue types. PL = polar lipids, NL = neutral lipids

| | | **df** | **Sum Sq.** | **Mean Sq.** | ***F*** | **Pr(>*F*)** |
|---|---|---|---|---|---|---|
| Brain PL | Ecotype | 2 | 8.9 | 4.4 | 2.93 | 0.081 |
| | Sex | 1 | 0.5 | 0.5 | 0.29 | 0.595 |
| | Residuals | 17 | 25.8 | 1.5 | | |
| Brain NL | Ecotype | 2 | 31.3 | 15.6 | 19.01 | < 0.001 |
| | Sex | 1 | < 0.1 | < 0.1 | 0.03 | 0.856 |
| | Residuals | 17 | 14.0 | 0.8 | | |
| Muscle PL | Ecotype | 2 | 10.1 | 5.1 | 2.32 | 0.129 |
| | Sex | 1 | 3.6 | 3.6 | 1.64 | 0.217 |
| | Residuals | 17 | 37.1 | 2.2 | | |
| Muscle NL | Ecotype | 2 | 11.6 | 5.8 | 4.58 | 0.026 |
| | Sex | 1 | 0.6 | 0.6 | 0.43 | 0.519 |
| | Residuals | 17 | 21.4 | 1.3 | | |

**Table S11.** Pairwise comparisons from Tukey's HSD *post hoc* tests of the effect of ecotype on ARA (20:4n-6) content (%) in neutral lipids (NL) in brain and muscle tissue.

| | **Comparison** | **Difference** | **Lower** | **Upper** | $p_{adj}$ |
|---|---|---|---|---|---|
| Brain NL | Pot : Ana | -0.788 | -2.031 | 0.456 | 0.262 |
| | Riv : Ana | -2.891 | -4.135 | -1.648 | < 0.001 |
| | Riv : Pot | -2.103 | -3.347 | -0.860 | 0.001 |
| Muscle NL | Pot : Ana | -0.580 | -2.121 | 0.960 | 0.607 |
| | Riv : Ana | -1.782 | -3.322 | -0.242 | 0.022 |
| | Riv : Pot | -1.201 | -2.741 | 0.339 | 0.142 |

**Table S12.** Pairwise comparisons from *post hoc* (Tukey's method) test of the effect of lipid class and tissue type on contents (%) of EPA (20:5n-3), DHA (22:6n-3) and ARA (20:4n-6). FAME = fatty acid methyl ester, PL = polar lipids, NL = neutral lipids

| FAME | Comparison | SE | df | *t* ratio | *p* value |
|---|---|---|---|---|---|
| EPA (20:5n-3) | brain NL : brain PL | -0.165 | 0.516 | -0.321 | 0.989 |
| | brain NL : muscle NL | -3.629 | 0.516 | -7.033 | < 0.001 |
| | brain NL : muscle PL | -2.060 | 0.516 | -3.992 | 0.001 |
| | brain PL : muscle NL | -3.463 | 0.516 | -6.712 | < 0.001 |
| | brain PL : muscle PL | -1.894 | 0.516 | -3.671 | 0.003 |
| | muscle NL : muscle PL | 1.569 | 0.516 | 3.041 | 0.018 |
| DHA (22:6n-3) | brain NL : brain PL | -23.18 | 1.2 | -19.24 | < 0.001 |
| | brain NL : muscle NL | -1.56 | 1.2 | -1.29 | 0.572 |
| | brain NL : muscle PL | -24.72 | 1.2 | -20.52 | < 0.001 |
| | brain PL : muscle NL | 21.63 | 1.2 | 17.95 | < 0.001 |
| | brain PL : muscle PL | -1.53 | 1.2 | -1.27 | 0.584 |
| | muscle NL : muscle PL | 23.16 | 1.2 | 19.22 | < 0.001 |
| ARA (20:4n-6) | brain NL : brain PL | -0.403 | 0.287 | -19.24 | < 0.001 |
| | brain NL : muscle NL | -1.083 | 0.287 | -1.29 | 0.572 |
| | brain NL : muscle PL | -2.977 | 0.287 | -20.52 | < 0.001 |
| | brain PL : muscle NL | -0.680 | 0.287 | 17.95 | < 0.001 |
| | brain PL : muscle PL | -2.574 | 0.287 | -1.27 | 0.584 |
| | muscle NL : muscle PL | -1.895 | 0.287 | 19.22 | < 0.001 |

* Degrees of freedom calculated using Kenward-Roger method

**Table S13.** ANOVA results for effects of ecotype and sex on $\Delta\delta^2H_{DHA}$ in two lipid classes from two tissue types. PL = polar lipids, NL = neutral lipids

| | | df | Sum Sq. | Mean Sq. | *F* | Pr(>*F*) |
|---|---|---|---|---|---|---|
| Brain PL | Ecotype | 2 | 111 | 56 | 0.53 | 0.599 |
| | Sex | 1 | 152 | 152 | 1.44 | 0.246 |
| | Residuals | 17 | 1790 | 105 | | |
| Brain NL | Ecotype | 2 | 695 | 347 | 0.92 | 0.423 |
| | Sex | 1 | 173 | 173 | 0.46 | 0.510 |
| | Residuals | 14 | 5312 | 380 | | |
| Muscle PL | Ecotype | 2 | 100 | 50 | 0.60 | 0.561 |
| | Sex | 1 | 1 | 1 | 0.01 | 0.937 |
| | Residuals | 16 | 1339 | 34 | | |
| Muscle NL | Ecotype | 2 | 568 | 284 | 0.97 | 0.402 |
| | Sex | 1 | 49 | 49 | 0.17 | 0.689 |
| | Residuals | 16 | 4713 | 295 | | |

**Table S14.** ANOVA results for effects of ecotype and sex on $\Delta\delta^2H_{ARA}$ in two lipid classes from two tissue types. PL = polar lipids, NL = neutral lipids

| | | **df** | **Sum Sq.** | **Mean Sq.** | ***F*** | **Pr(>*F*)** |
|---|---|---|---|---|---|---|
| Brain PL | Ecotype | 2 | 10077 | 5038 | 5.464 | 0.018 |
| | Sex | 1 | 407 | 407 | 0.442 | 0.517 |
| | Residuals | 14 | 12909 | 922 | | |
| Brain NL | Ecotype | 2 | 2081 | 1041 | 1.906 | 0.199 |
| | Sex | 1 | 2 | 2 | 0.005 | 0.948 |
| | Residuals | 10 | 5460 | 546 | | |
| Muscle PL | Ecotype | 2 | 22339 | 11169 | 15.05 | < 0.001 |
| | Sex | 1 | 30 | 30 | 0.04 | 0.844 |
| | Residuals | 14 | 10387 | 742 | | |
| Muscle NL | Ecotype | 2 | 4899 | 2449 | 5.078 | 0.020 |
| | Sex | 1 | 230 | 230 | 0.476 | 0.500 |
| | Residuals | 16 | 7718 | 482 | | |

**Table S15.** Pairwise comparisons from Tukey's HSD *post hoc* tests of the effect of ecotype on $\Delta\delta^2H_{ARA}$ in brain and muscle lipids. PL = polar lipids, NL = neutral lipids

| | Comparison | Difference | Lower | Upper | $p$ adj |
|---|---|---|---|---|---|
| Brain PL | Pot : Ana | -5.14 | -47.62 | 37.34 | 0.946 |
| | Riv : Ana | -59.22 | -109.03 | -9.41 | 0.020 |
| | Riv : Pot | -54.08 | -103.90 | -4.27 | 0.033 |
| Muscle PL | Pot : Ana | 25.94 | -13.73 | 65.60 | 0.236 |
| | Riv : Ana | -62.76 | -104.50 | -21.01 | 0.004 |
| | Riv : Pot | -88.69 | -131.86 | -45.53 | < 0.001 |
| Muscle NL | Pot : Ana | -5.84 | -37.37 | 25.69 | 0.883 |
| | Riv : Ana | -35.14 | -65.43 | -4.84 | 0.022 |
| | Riv : Pot | -29.23 | -60.83 | -2.23 | 0.071 |

**Table S16.** Probabilities of individual brown trout being anadromous, potamodromous and riverine resident, determined by a multinomial logistic regression of ecotype on DHA and EPA contents (%) in polar and neutral lipids in brain and muscle tissue.

| | | **Probabilities** | | |
|---|---|---|---|---|
| **ID** | **Actual type** | **Anadromous** | **Potamodromous** | **Riverine** |
| E07 | Anadromous | 1.00 | 0.00 | 0.00 |
| E12 | Anadromous | 1.00 | 0.00 | 0.00 |
| E21 | Anadromous | 1.00 | 0.00 | 0.00 |
| E26 | Anadromous | 1.00 | 0.00 | 0.00 |
| E33 | Anadromous | 1.00 | 0.00 | 0.00 |
| E35 | Anadromous | 1.00 | 0.00 | 0.00 |
| E43 | Anadromous | 1.00 | 0.00 | 0.00 |
| E01 | Potamodromous | 0.00 | 1.00 | 0.00 |
| E02 | Potamodromous | 0.00 | 1.00 | 0.00 |
| E06 | Potamodromous | 0.00 | 1.00 | 0.00 |
| E42 | Potamodromous | 0.00 | 1.00 | 0.00 |
| E46 | Potamodromous | 0.00 | 1.00 | 0.00 |
| E54 | Potamodromous | 0.00 | 1.00 | 0.00 |
| E58 | Potamodromous | 0.00 | 1.00 | 0.00 |
| E61 | Riverine | 0.00 | 0.00 | 1.00 |
| E63 | Riverine | 0.00 | 0.00 | 1.00 |
| E64 | Riverine | 0.00 | 0.00 | 1.00 |
| E65 | Riverine | 0.00 | 0.00 | 1.00 |
| E66 | Riverine | 0.00 | 0.00 | 1.00 |
| E69 | Riverine | 0.00 | 0.00 | 1.00 |
| E70 | Riverine | 0.00 | 0.00 | 1.00 |

**Table S17.** Probabilities of individual brown trout being anadromous, potamodromous and riverine resident, determined by a multinomial logistic regression of ecotype on DHA and ARA content (%) in polar and neutral lipids in brain and muscle tissue.

| | | Probabilities | | |
|---|---|---|---|---|
| **ID** | **Actual type** | **Anadromous** | **Potamodromous** | **Riverine** |
| E07 | Anadromous | 1.00 | 0.00 | 0.00 |
| E12 | Anadromous | 1.00 | 0.00 | 0.00 |
| E21 | Anadromous | 1.00 | 0.00 | 0.00 |
| E26 | Anadromous | 0.99 | 0.01 | 0.00 |
| E33 | Anadromous | 0.99 | 0.01 | 0.00 |
| E35 | Anadromous | 1.00 | 0.00 | 0.00 |
| E43 | Anadromous | 0.92 | 0.08 | 0.00 |
| E01 | Potamodromous | 0.00 | 1.00 | 0.00 |
| E02 | Potamodromous | 0.00 | 1.00 | 0.00 |
| E06 | Potamodromous | 0.02 | 0.98 | 0.00 |
| E42 | Potamodromous | 0.02 | 0.98 | 0.00 |
| E46 | Potamodromous | 0.02 | 0.98 | 0.00 |
| E54 | Potamodromous | 0.03 | 0.97 | 0.00 |
| E58 | Potamodromous | 0.00 | 1.00 | 0.00 |
| E61 | Riverine | 0.00 | 0.00 | 1.00 |
| E63 | Riverine | 0.00 | 0.00 | 1.00 |
| E64 | Riverine | 0.00 | 0.00 | 1.00 |
| E65 | Riverine | 0.00 | 0.00 | 1.00 |
| E66 | Riverine | 0.00 | 0.00 | 1.00 |
| E69 | Riverine | 0.00 | 0.00 | 1.00 |
| E70 | Riverine | 0.00 | 0.00 | 1.00 |

**Table S18.** Probabilities of individual brown trout being anadromous, potamodromous and riverine resident, determined by a multinomial logistic regression of ecotype on $\Delta\delta^2H$ of DHA and EPA in polar and neutral lipids in brain and muscle tissue.

| | | Probabilities | | |
|---|---|---|---|---|
| **ID** | **Actual type** | **Anadromous** | **Potamodromous** | **Riverine** |
| E07 | Anadromous | 1.00 | 0.00 | 0.00 |
| E12 | Anadromous | 1.00 | 0.00 | 0.00 |
| E21 | Anadromous | 1.00 | 0.00 | 0.00 |
| E26 | Anadromous | 1.00 | 0.00 | 0.00 |
| E33 | Anadromous | 1.00 | 0.00 | 0.00 |
| E35 | Anadromous | 1.00 | 0.00 | 0.00 |
| E01 | Potamodromous | 0.00 | 1.00 | 0.00 |
| E06 | Potamodromous | 0.00 | 1.00 | 0.00 |
| E46 | Potamodromous | 0.00 | 1.00 | 0.00 |
| E54 | Potamodromous | 0.00 | 1.00 | 0.00 |
| E58 | Potamodromous | 0.00 | 1.00 | 0.00 |
| E61 | Riverine | 0.00 | 0.00 | 1.00 |
| E64 | Riverine | 0.00 | 0.00 | 1.00 |
| E65 | Riverine | 0.00 | 0.00 | 1.00 |
| E66 | Riverine | 0.00 | 0.00 | 1.00 |
| E69 | Riverine | 0.00 | 0.00 | 1.00 |

**Table S19.** Probabilities of individual brown trout being anadromous, potamodromous and riverine resident, determined by a multinomial logistic regression of ecotype on $\Delta\delta^2H$ of DHA and ARA in polar and neutral lipids in brain and muscle tissue.

| | | Probabilities | | |
|---|---|---|---|---|
| **ID** | **Actual type** | **Anadromous** | **Potamodromous** | **Riverine** |
| E07 | Anadromous | 1.00 | 0.00 | 0.00 |
| E12 | Anadromous | 1.00 | 0.00 | 0.00 |
| E21 | Anadromous | 1.00 | 0.00 | 0.00 |
| E26 | Anadromous | 1.00 | 0.00 | 0.00 |
| E33 | Anadromous | 1.00 | 0.00 | 0.00 |
| E35 | Anadromous | 1.00 | 0.00 | 0.00 |
| E01 | Potamodromous | 0.00 | 1.00 | 0.00 |
| E46 | Potamodromous | 0.00 | 1.00 | 0.00 |
| E54 | Potamodromous | 0.00 | 1.00 | 0.00 |
| E65 | Riverine | 0.00 | 0.00 | 1.00 |
| E66 | Riverine | 0.00 | 0.00 | 1.00 |

**Table S21.** Results from ANOVA testing the association of brown trout migratory ecotype with gene co-expression module, with Tukey HSD *post-hoc* results where appropriate. A = anadromous, P = potamodromous, R = riverine.

| **ANOVA** | | | | **Tukey HSD** | | | | |
|---|---|---|---|---|---|---|---|---|
| **Module** | ***F* value** | ***p* value** | ***q* value** | **Comparison** | **Difference** | **Lower** | **Upper** | ***p* value (adj.)** |
| Coral3 | 3.752 | 0.043 | 0.064 | P-A | 2.429 | -5.067 | 9.924 | 0.692 |
| | | | | R-A | 7.857 | 0.361 | 15.353 | 0.039 |
| | | | | R-P | 5.429 | -2.067 | 12.924 | 0.183 |
| Palevioletred2 | 4.672 | 0.023 | 0.050 | P-A | 7.286 | 0.047 | 14.525 | 0.048 |
| | | | | R-A | 7.714 | 0.475 | 14.953 | 0.036 |
| | | | | R-P | 0.429 | -6.810 | 7.668 | 0.988 |
| Yellow4 | 4.386 | 0.028 | 0.052 | P-A | 7.857 | 0.541 | 15.173 | 0.034 |
| | | | | R-A | 6.714 | -0.602 | 14.030 | 0.075 |
| | | | | R-P | -1.143 | -8.459 | 6.173 | 0.917 |
| Darkorange2 | 4.019 | 0.036 | 0.059 | *n.a.* | | | | |
| Black | 18.131 | < 0.001 | < 0.001 | P-A | -0.143 | -5.282 | 4.996 | 0.997 |
| | | | | R-A | 10.429 | 5.290 | 15.568 | < 0.001 |
| | | | | R-P | 10.571 | 5.432 | 15.710 | < 0.001 |
| Firebrick3 | 7.366 | 0.005 | 0.013 | P-A | 2.429 | -4.188 | 9.045 | 0.625 |
| | | | | R-A | 9.571 | 2.955 | 16.188 | 0.005 |
| | | | | R-P | 7.143 | 0.526 | 13.759 | 0.033 |
| Antiquewhite1 | 8.691 | 0.002 | 0.007 | P-A | 0.286 | -6.078 | 6.650 | 0.993 |
| | | | | R-A | 9.143 | 2.779 | 15.507 | 0.005 |
| | | | | R-P | 8.857 | 2.493 | 15.221 | 0.006 |
| Lavenderblush2 | 3.874 | 0.040 | 0.062 | *n.a.* | | | | |
| Skyblue2 | 14.166 | < 0.001 | 0.001 | P-A | 4.000 | -1.561 | 9.561 | 0.187 |
| | | | | R-A | 11.429 | 5.867 | 16.990 | < 0.001 |
| | | | | R-P | 7.429 | 1.867 | 12.990 | 0.008 |
| Floralwhite | 0.884 | 0.430 | 0.482 | *n.a.* | | | | |
| Skyblue | 0.208 | 0.814 | 0.839 | *n.a.* | | | | |
| Green4 | 0.290 | 0.752 | 0.810 | *n.a.* | | | | |
| Mediumpurple1 | 1.286 | 0.301 | 0.351 | *n.a.* | | | | |
| Pink | 0.177 | 0.839 | 0.839 | *n.a.* | | | | |
| Lightsteelblue1 | 4.430 | 0.027 | 0.052 | *n.a.* | | | | |
| Blueviolet | 5.209 | 0.016 | 0.038 | P-A | -5.714 | -12.815 | 1.387 | 0.128 |
| | | | | R-A | -8.857 | -15.958 | -1.756 | 0.014 |
| | | | | R-P | -3.143 | -10.244 | 3.958 | 0.509 |
| Coral | 14.233 | < 0.001 | 0.001 | P-A | -3.286 | -8.839 | 2.268 | 0.310 |
| | | | | R-A | -11.286 | -16.839 | -5.732 | < 0.001 |
| | | | | R-P | -8.000 | -13.553 | -2.447 | 0.005 |
| Ivory | 6.361 | 0.008 | 0.021 | P-A | -6.000 | -12.830 | 0.830 | 0.091 |
| | | | | R-A | -9.429 | -16.258 | -2.599 | 0.007 |
| | | | | R-P | -3.429 | -10.258 | 3.401 | 0.423 |
| Chocolate4 | 8.679 | 0.002 | 0.007 | P-A | 0.000 | -6.366 | 6.366 | 1.000 |
| | | | | R-A | -9.000 | -15.366 | -2.634 | 0.005 |
| | | | | R-P | -9.000 | -15.366 | -2.634 | 0.005 |
| Lightsteelblue | 14.503 | < 0.001 | 0.001 | P-A | -0.714 | -6.236 | 4.807 | 0.942 |
| | | | | R-A | -10.429 | -15.950 | -4.907 | < 0.001 |
| | | | | R-P | -9.714 | -15.236 | -4.193 | < 0.001 |
| Blue | 8.679 | 0.002 | 0.007 | P-A | 0.000 | -6.366 | 6.366 | 1.000 |
| | | | | R-A | -9.000 | -15.366 | -2.634 | 0.005 |
| | | | | R-P | -9.000 | -15.366 | -2.634 | 0.005 |
| Steelblue | 4.276 | 0.030 | 0.053 | *n.a.* | | | | |
| Magenta3 | 3.413 | 0.055 | 0.078 | *n.a.* | | | | |
| Darkslateblue | 2.972 | 0.077 | 0.102 | P-A | 2.571 | -5.165 | 10.308 | 0.679 |
| | | | | R-A | -4.714 | -12.450 | 3.022 | 0.290 |
| | | | | R-P | -7.286 | -15.022 | 0.450 | 0.067 |
| Antiquewhite2 | 18.752 | < 0.001 | < 0.001 | P-A | 1.286 | -3.795 | 6.367 | 0.797 |
| | | | | R-A | -9.857 | -14.938 | -4.776 | < 0.001 |
| | | | | R-P | -11.143 | -16.224 | -6.062 | < 0.001 |
| Darkseagreen2 | 11.314 | < 0.001 | 0.003 | P-A | 2.000 | -3.939 | 7.939 | 0.672 |
| | | | | R-A | -8.429 | -14.367 | -2.490 | 0.005 |
| | | | | R-P | -10.429 | -16.367 | -4.490 | < 0.001 |
| Plum2 | 2.872 | 0.083 | 0.105 | *n.a.* | | | | |
| Thistle2 | 1.737 | 0.204 | 0.249 | *n.a.* | | | | |

**Table S22.** Gene ontology terms for genes within the 'black' co-expression module.

| **Term name** | **Term ID** | **Highlighted** | ***p* adj.** |
|---|---|---|---|
| Cell communication | GO:0007154 | True | < 0.001 |
| Response to stimulus | GO:0050896 | False | < 0.001 |
| Signalling | GO:0023052 | False | < 0.001 |
| Signal transduction | GO:0007165 | False | < 0.001 |
| Cellular response to stimulus | GO:0051716 | False | < 0.001 |
| Immune system process | GO:0002376 | True | < 0.001 |
| Cell cycle process | GO:0022402 | True | 0.002 |
| Mitotic cell cycle process | GO:1903047 | False | 0.012 |
| Regulation of biological process | GO:0050789 | False | 0.023 |
| Organelle organisation | GO:0006996 | True | 0.027 |
| Chromosome organisation | GO:0051276 | False | 0.028 |
| Regulation of cellular process | GO:0050794 | False | 0.032 |

**Table S23.** Gene ontology terms for genes within the 'blue' co-expression module.

| Term name | Term ID | Highlighted | *p* adj. |
|---|---|---|---|
| Translation | GO:0006412 | True | < 0.001 |
| Peptide biosynthetic process | GO:0043043 | False | < 0.001 |
| Amide biosynthetic process | GO:0043604 | False | < 0.001 |
| Peptide metabolic process | GO:0006518 | False | < 0.001 |
| Amide metabolic process | GO:0043603 | False | < 0.001 |
| Organonitrogen compound biosynthetic process | GO:1901566 | False | < 0.001 |
| Cellular biosynthetic process | GO:0044249 | False | < 0.001 |
| Cellular nitrogen compound Biosynthetic process | GO:0044271 | False | < 0.001 |
| Macromolecule biosynthetic process | GO:0009059 | False | < 0.001 |
| Biosynthetic process | GO:0009058 | False | < 0.001 |
| Organic substance biosynthetic process | GO:1901576 | False | < 0.001 |
| Gene expression | GO:0010467 | False | < 0.001 |
| Cellular nitrogen compound metabolic process | GO:0034641 | False | < 0.001 |
| Cellular nitrogen compound metabolic process | GO:1901564 | False | < 0.001 |
| Protein metabolic process | GO:0019538 | False | < 0.001 |
| Cellular process | GO:0009987 | False | < 0.001 |
| Cellular metabolic process | GO:0044237 | False | < 0.001 |
| Biological process | GO:0008150 | False | < 0.001 |
| Organic substance metabolic process | GO:0071704 | False | < 0.001 |
| Nitrogen substance metabolic process | GO:0006807 | False | < 0.001 |
| Primary metabolic process | GO:0044238 | False | < 0.001 |
| Metabolic process | GO:0044238 | False | < 0.001 |
| Macromolecule metabolic process | GO:0043170 | False | < 0.001 |
| Translational elongation | GO:0006414 | False | < 0.001 |

**Table S24.** ANOVA results for effects of ecotype and sex on $\Delta\delta^2H_{EPA}$ in two lipid classes from two tissue types. PL = polar lipids, NL = neutral lipids

| | | **df** | **Sum Sq.** | **Mean Sq.** | ***F*** | **Pr(>*F*)** |
|---|---|---|---|---|---|---|
| Brain PL | Ecotype | 2 | 5592 | 2796 | 4.30 | 0.031 |
| | Sex | 1 | 13 | 13 | 0.02 | 0.891 |
| | Residuals | 17 | 11046 | 650 | | |
| Brain NL | Ecotype | 2 | 7205 | 3602 | 4.07 | 0.041 |
| | Sex | 1 | 122 | 122 | 0.14 | 0.717 |
| | Residuals | 14 | 12399 | 886 | | |
| Muscle PL | Ecotype | 2 | 3490 | 1745 | 3.36 | 0.061 |
| | Sex | 1 | 21 | 21 | 0.04 | 0.843 |
| | Residuals | 16 | 8312 | 519 | | |
| Muscle NL | Ecotype | 2 | 648 | 324 | 0.58 | 0.571 |
| | Sex | 1 | 247 | 3.1247 | 0.44 | 0.515 |
| | Residuals | 16 | 8925 | 558 | | |

**Table S25.** Pairwise comparisons from Tukey's HSD *post hoc* tests of the effect of ecotype on $\Delta\delta^2H_{EPA}$ in brain lipids. PL = polar lipids, NL = neutral lipids

| | **Comparison** | **Difference** | **Lower** | **Upper** | ***p* adj** |
|---|---|---|---|---|---|
| Brain PL | Pot : Ana | 6.556 | -28.397 | 41.509 | 0.881 |
| | Riv : Ana | 37.425 | 2.472 | 72.378 | 0.035 |
| | Riv : Pot | 30.869 | -4.084 | 65.822 | 0.089 |
| Brain NL | Pot : Ana | 10.742 | -32.592 | 54.076 | 0.796 |
| | Riv : Ana | 49.281 | 2.116 | 96.446 | 0.040 |
| | Riv : Pot | 38.529 | -7.069 | 84.147 | 0.104 |

**Table S26.** Results of Procrustes ANOVA testing effects of centroid size and ecotype on brown trout body shape, using a randomised residual permutation procedure with 1,000 permutations.

| | df | SS | MS | $R^2$ | *F* | *Z* | *p* |
|---|---|---|---|---|---|---|---|
| **Centroid size** | 1 | 0.0084 | 0.0084 | 0.130 | 17.35 | 4.97 | 0.001 |
| **Ecotype** | 2 | 0.0168 | 0.0084 | 0.261 | 17.36 | 6.01 | 0.001 |
| **Residuals** | 81 | 0.0392 | 0.0048 | 0.609 | | | |
| **Total** | 84 | 0.0644 | | | | | |

**Table S27.** Pairwise comparisons by ecotype of least-squares means distances. Ana = anadromous, Pot = potamodromous, Riv = riverine.

| | **distance** | **UCL (95%)** | ***Z*** | ***p*** |
|---|---|---|---|---|
| **Ana : Pot** | 0.0149 | 0.0105 | 2.965 | 0.001 |
| **Ana : Riv** | 0.0308 | 0.0116 | 4.969 | 0.001 |
| **Pot : Riv** | 0.0318 | 0.0103 | 5.003 | 0.001 |

**Table S28.** Primer sequences for PCR genotyping for sex of brown trout.

| **Primer** | **Sequence** |
|---|---|
| *sdY* forward | CCC AGC ACT GTT TTC TTG TCT CA |
| *sdY* reverse | CTT AAA ACC ACT CCA CCC TCC AT |
| *18S* forward | GTA CGA AGA CGA TCA GAT ACC GT |
| *18S* reverse | CCG CAT AAC TAG TTA GCA TGC CG |

**Table S29.** Duplex PCR mix (per sample) for sex genotyping of brown trout.

| **Reagent** | **Volume** | |
|---|---|---|
| Qiagen Multiplex PCR mix * | 7.5 | µL |
| *sdY* forward primer | 0.3 | µL |
| *sdY* reverse primer | 0.3 | µL |
| *18S* forward primer | 0.075 | µL |
| *18S* reverse primer | 0.075 | µL |
| $H_2O$ (nuclease-free) | 2.25 | µL |
| DNA (20 ng $\mu L^{-1}$) | 3 | µL |

* containing 3 mM $MgCl_2$, HotStarTaq DNA polymerase and proprietary buffer

**Table S30.** Thermocycling protocol for sex genotyping of brown trout.

| Stage | Temperature | Duration | Cycles |
|---|---|---|---|
| Initialisation | 95 °C | 15 min | 1 |
| Amplification | 94 °C | 30 sec | 35 |
| | 63 °C | 90 sec | |
| | 72 °C | 90 sec | |
| Final extension | 72 °C | 10 min | 1 |

**Table S31.** Brown trout liver samples used for mRNAseq analyses, corresponding to compound-specific stable isotope analyses of brain and muscle tissue, with ecotype and sex designations, fork lengths (FL in mm), and number of mRNA sequence read counts, including those assigned to no feature (*i.e.* gene), or were ambiguous or not unique.

| | | | | mRNA sequence read counts | | | |
|---|---|---|---|---|---|---|---|
| **ID** | **Ecotype** | **Sex** | **FL** | **Total** | **No feature** | **Ambiguous** | **Not unique** |
| E07 | Anadromous | Female | 131 | 42,402,790 | 4,756,017 | 902,652 | 4,781,576 |
| E12 | Anadromous | Female | 127 | 23,028,156 | 2,330,885 | 564,252 | 2,393,293 |
| E21 | Anadromous | Male | 121 | 54,704,170 | 5,883,608 | 1,301,022 | 5,610,005 |
| E26 | Anadromous | Male | 126 | 21,847,686 | 2,267,424 | 591,887 | 2,778,502 |
| E33 | Anadromous | Male | 145 | 39,884,554 | 4,336,799 | 1,412,313 | 4,564,771 |
| E35 | Anadromous | Female | 139 | 38,972,830 | 4,538,967 | 845,113 | 3,930,750 |
| E43 | Anadromous | Male | 145 | 39,329,166 | 4,808,309 | 933,566 | 4,227,966 |
| E01 | Potamodromous | Female | 110 | 41,813,980 | 4,476,878 | 847,615 | 4,095,983 |
| E02 | Potamodromous | Male | 113 | 23,517,846 | 2,708,878 | 534,527 | 2,555,269 |
| E06 | Potamodromous | Female | 105 | 37,867,157 | 3,604,875 | 797,931 | 3,879,933 |
| E42 | Potamodromous | Male | 120 | 39,707,961 | 3,509,607 | 975,809 | 3,588,177 |
| E46 | Potamodromous | Female | 114 | 40,080,840 | 4,671,744 | 789,670 | 4,943,540 |
| E54 | Potamodromous | Male | 131 | 43,987,505 | 4,890,559 | 1,223,593 | 4,742,388 |
| E58 | Potamodromous | Male | 109 | 41,322,326 | 4,043,482 | 971,290 | 4,558,260 |
| E61 | Riverine | Male | 67 | 35,547,150 | 3,274,803 | 789,764 | 4,668,912 |
| E63 | Riverine | Male | 68 | 50,909,690 | 5,338,898 | 1,130,600 | 5,663,094 |
| E64 | Riverine | Male | 65 | 41,256,877 | 3,562,676 | 944,980 | 5,346,975 |
| E65 | Riverine | Male | 61 | 37,598,658 | 3,957,311 | 847,193 | 4,600,188 |
| E66 | Riverine | Female | 79 | 42,857,777 | 4,362,624 | 1,092,557 | 5,344,053 |
| E69 | Riverine | Female | 68 | 31,822,339 | 3,005,561 | 757,481 | 4,338,902 |
| E70 | Riverine | Female | 70 | 21,124,240 | 2,005,584 | 444,664 | 2,643,713 |
